# Binding Profile Mapping of the S100 Protein Family Using a High-throughput Local Surface Mimetic Holdup Assay

**DOI:** 10.1101/2020.12.02.407676

**Authors:** Márton A. Simon, Éva Bartus, Beáta Mag, Eszter Boros, Lea Roszjár, Gergő Gógl, Gilles Travé, Tamás A. Martinek, László Nyitray

## Abstract

S100 proteins are small, typically homodimeric, vertebrate-specific EF-hand proteins that establish Ca^2+^-dependent protein-protein interactions in the intra- and extracellular environment and are overexpressed in various pathologies. There are about 20 distinct human S100 proteins with numerous potential partner proteins. Here, we used a quantitative holdup assay to measure affinity profiles of most members of the S100 protein family against a library of chemically synthetized foldamers. The profiles allowed us to quantitatively map the binding promiscuity of each member towards the foldamer library. Since the library was designed to systematically contain most binary natural amino acid side chain combinations, the data also provide insight into the promiscuity of each S100 protein towards all potential naturally-occurring S100 partners in the human proteome. Such information will be precious for future drug design of modulators of S100 pathological activities.

## Introduction

The vertebrate-specific calcium-binding S100 protein family (termed here as the S100ome) belongs to the superfamily of the EF-hand containing proteins and consists of at least 20 core members of small (10 kDa), usually homodimeric proteins that play role in cellular regulation both intra- and extracellularly via protein-protein interactions (PPIs) in a Ca^2+^-dependent manner [1, 2]. Under physiological conditions, their expression pattern is tissue-specific and they are present usually in low concentrations. However, their expression level and pattern can be altered under pathological conditions, leading to severe consequences [3]. Specifically, elevated cellular concentrations of certain S100 proteins were observed in cancer, cardiomyopathies, inflammatory and neurodegenerative diseases [3, 4], pointing to them as potential biomarkers and/or therapeutic targets of these diseases [5]. Development of selective inhibitors have great pharmaceutical potential, but it is still challenging due to the structural similarity within the S100 family. Thus, comprehensive and accurate mapping of the specific S100 interactome is required for such purpose [6]. Although numerous S100 binding partners are known, they are rather restricted to a small subset of the protein family (e.g. S100B, S100A4) [1]. Therefore, a family-wide systematic screening is in need to map the specificity and affinity profiles within the entire S100 family and to identify new binding partners.

Experimental characterization of protein surfaces having shallow binding clefts is a great challenge in drug discovery; however, tools of fragment-based approaches have become efficient techniques toward identification of small-molecule drug candidates [7]. Mapping the binding surface of proteins can be performed with short recognition elements (i.e. small patches of the binding interface) displaying reduced structural complexity [8–10]. Local surface mimetic (LSM) foldamers having water-stable H14 helical conformation proved to be efficient probes for screening these shallow binding clefts of different protein targets [11]. In a high-throughput (HTP) experimental setup, the binding properties of the S100ome can be characterized comprehensively by using this LSM foldamer library. The library members are able to point two proteinogenic side chains at *i* and *i+3* positions toward the surface and thereby mimic the small pieces of the complementary binding interface for the tested protein. A 256-member foldamer library was designed to cover all the possible combination pairs of 16 different proteinogenic side chains [12]. Regardless of their secondary structure, LSMs can mimic binary surface fragments with their 5 Å distances between the proteinogenic side chains resulting small surface patches rather than secondary structure mimetics. (**Fig 1A**). The bulky cyclic amino acids in the helix (trans-2-aminocyclohexancarboxylic acid) shield the proteinogenic side chains introduced by β^3^-amino acid derivatives from the solvents [11].

**Fig. 1.**
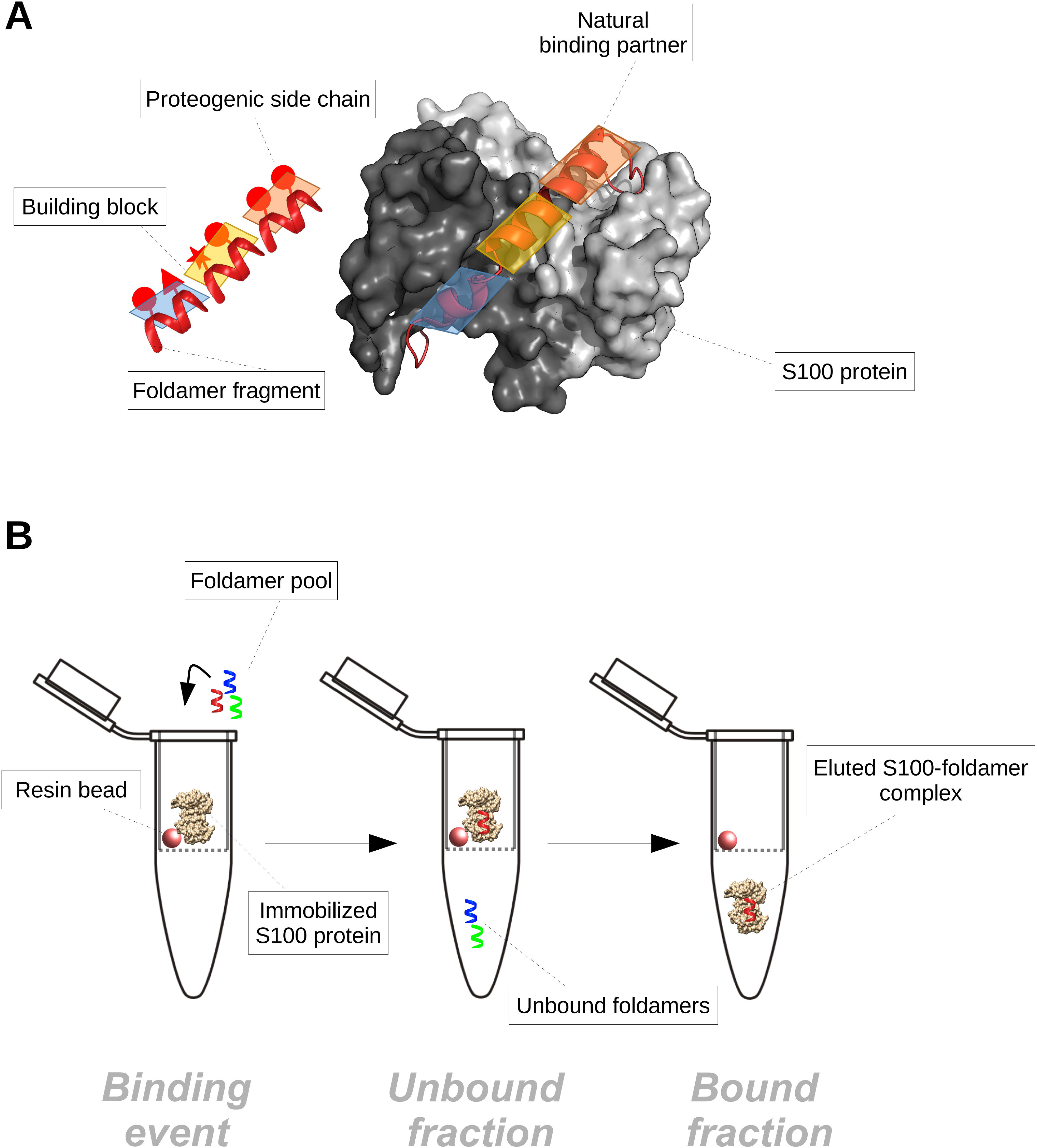
The methodology of the high-throughput (HTP) holdup (HU) assay. Panel A: S100 proteins generally prefer helical partners or disordered partners prone to helix formation upon binding (right: S100A4 – NMIIA complex [29], PDB code: 3ZWH). Foldamer fragments (left: H14 foldamer building blocks, each with two proteinogenic side chain) cannot act as perfect α-helical mimetics, as the introduction of β-amino acids results in a bulkier helix with an altered amino acid pattern compared to the α-helix. They rather act as local surface patches, which present the proteogenic amino acids to the complementary binding surface; while the hydrophobic frame exclude the solvent from the binding pocket. The S100 binding surface can accommodate multiple foldamer fragments. Panel B: His-tagged S100 proteins immobilized on Co^2+^-resin (left panel) are incubated with the H14 foldamer library (256 members). The unbound fraction (flow-through) is recovered (middle panel) and the resin is submitted to a washing step (not represented). Finally, the S100 protein of interest together with the bound foldamer fragments is eluted by adding imidazole (right panel). Both the flow-through and eluted fractions are analyzed by LC-MS.

The members of the S100 family are often regarded as rather unspecific, promiscuous proteins [13]. Based on our recent study, the S100ome can be divided into two groups, according to binding preference against several natural S100 partners [6]. The partner preferences give a good approximation for the classification of S100 member with multiple partners; nevertheless, the specificity and affinity profile of S100 proteins without a clear binding preference (orphan) are still unknown. Here we reasoned that the binding surface of the S100ome could be mapped extensively by the application of the foldamer-based library containing most natural side chain combination, which cover the general side chain preference of the S100ome by mimicking the complementary binding surface of interacting partners. In this study, we thoroughly investigated the general and unique characteristics of the binding surface of the S100ome by determining the binding affinities of the diverse H14 LSM foldamer library towards the S100 proteins in a HTP holdup (HU) assay [12, 14, 15]. Our experimental results revealed the binding preferences of not only S100 proteins with multiple known interactions but also S100 members lacking known interaction partners (orphans) in living organisms.

## Results

### Screening the binding affinities of the S100ome against the LSM library by a HTP HU assay

We screened the binding affinities of the S100ome towards the LSM library by using a HTP HU assay (**Fig 1B**), in which the 256-member LSM library was divided into four sub-libraries (each containing 64 individual foldamer fragments). S100 proteins were immobilized on Co^2+^-resin through their N-terminal His_6_-tag, and incubated with the foldamer sublibraries. Experimental conditions were set so that each S100 protein was in equimolar amount (64 μM) with the global concentration of the foldamer sub-library (containing the 64 foldamer fragments in 1 μM), thus all foldameric fragments had the opportunity to bind to the protein target, as described previously [12]. After the co-incubation, the unbound foldamers (the flow-through fraction) were separated from the protein-foldamer complexes (resin-bound fraction). Samples were analyzed on LC-MS system, and library members were characterized quantitatively in all samples by their area under the curve (AUC) in the total ion chromatograms. The AUC value of the appropriate foldameric element in the flow-through fraction was compared to a control sample (comprising all the components of the assay except the immobilized S100 protein) prepared under the same conditions. In this way, we quantified the fraction of each foldamer that was specifically retained on the resin containing immobilized S100 protein. This approach allowed us to determine a bound fraction (F_b_) which we used to calculate apparent K_d_ values.

We used this approach to map the binding affinities of the complete S100ome and determined the apparent dissociation constants of 5120 interactions (20 S100 proteins versus the 256-member foldamer library), depicted as heat maps (**Fig 2A, Fig S1**). The binding patterns of the S100 proteins for LSMs were found to be highly diverse. Some S100 members (e.g. S100A16, S100G) displayed only weak interactions (K_d_ > 100μM) toward the foldamers, while other family members (e.g. S100A2, S100A6) showed high propensity to bind LSM probes (**Fig S1**). The pair of identical hydrophobic binding pockets in S100 homodimers [2], created by the Ca^2+^-induced conformational changes, could be recognized by highly hydrophobic side chains with limited selectivity and the LSM library generally displayed enrichment for residues Trp, Phe, Ile and Leu. Beside the most favored hydrophobic side chains, which can often be observed in systematic libraries [16], foldamers containing basic and polar residues were also enriched on the protein binding sites in some cases, providing useful information to increase selectivity in rational drug design (**Fig 2B**). S100 family members are rather acidic proteins (pI_average_ = 5.68 ± 0.92), therefore, basic residues (Arg and Lys) are preferred in their ligands over acidic side chains (Glu and Asp). The enrichments of positively charged residues were found significant in our assay for S100A1, S100A2, S100B and S100P; as these S100 family members possess the lowest theoretical pIs (4.39, 4.68, 4.52 and 4.75, respectively). It is notable that neutral polar side chains were also found preferable for some of these family members (e.g. S100A2, S100P).

**Fig. 2.**
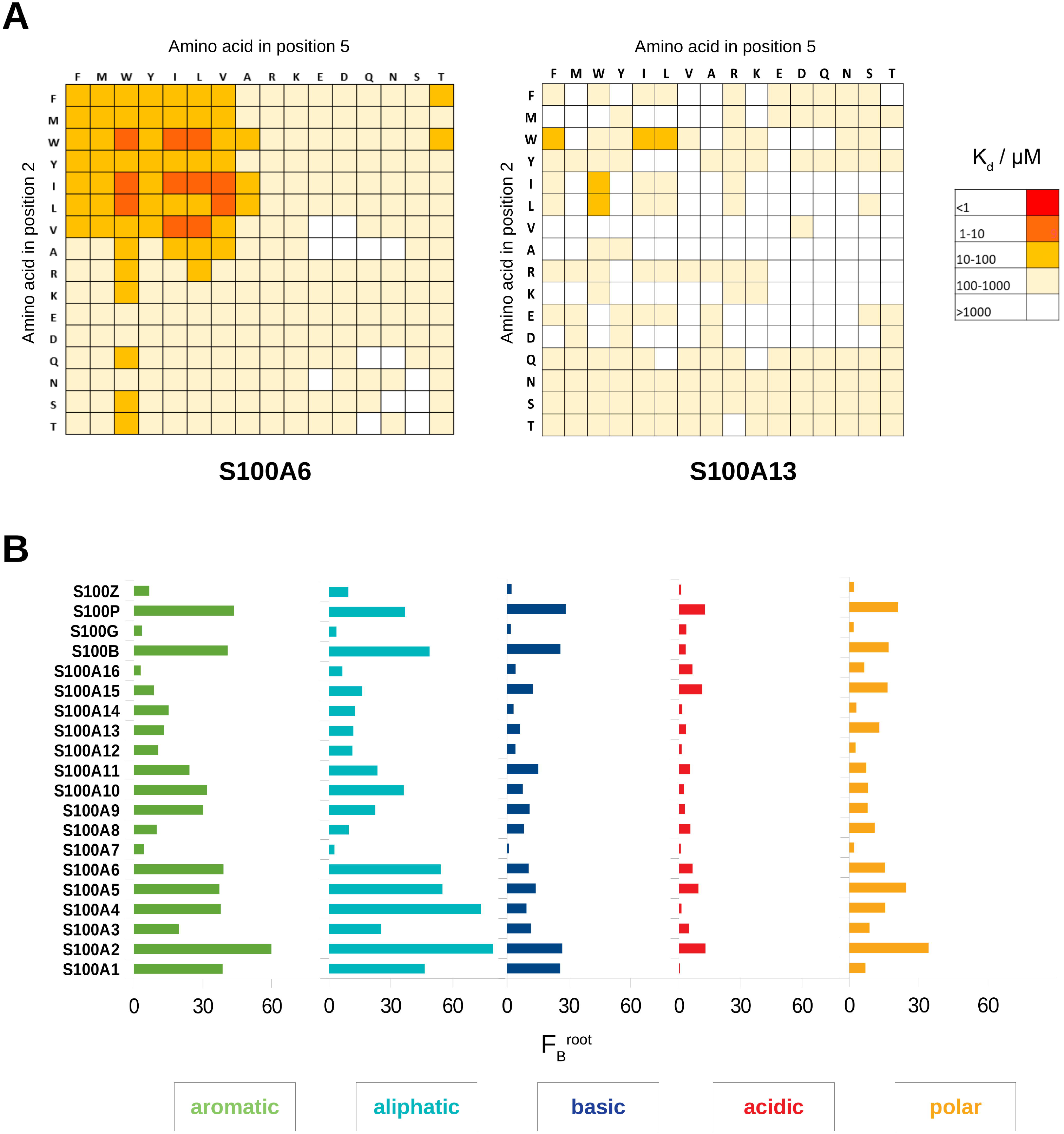
The interaction between the H14 foldamer library and the S100ome measured by holdup (HU) assay. Panel A: The interactions between S100 proteins and foldamers were measured by a high-throughput (HTP) holdup (HU) assay, as visualized in **Fig 1B**. Dissociation constants were calculated based on the loss of intensity of the foldamer of interest in the flow-through fraction using **eq. 1** and **eq. 2**(see materials and methods); and were depicted as a heat map in linear scale for each S100 protein. K_d_ ranges are color coded as shown on the right. The vertical axis and horizontal axis represents the β-amino acid in the second and fifth positions, respectively [11]. Some S100 members favored multiple fragments (e.g. S100A6 on the left), while multiple S100 proteins did not show clear binding preference towards the foldamer fragments (e.g. S100A13 on the right). Panel B: S100 proteins exert different amino acid sidechain preference based on the HTP HU measurements. The amino acid preferences were calculated for all S100 proteins using **eq. 4** and **eq. 5**(see materials and methods), and F_B_^root^ values were depicted as a bar chart. The residues with high frequency in the bound foldamers have hydrophobic properties as aromatic and aliphatic side chains are the most preferred ones. Importantly, due to the rather acidic nature of S100 proteins, acidic side chains are the least preferred among S100 proteins. It is noteworthy that in some instances polar residues are also favored (e.g. S100A2, S100A5).

The binding pattern mostly displayed a diagonal symmetry indicating a neutral template nature of the backbone. For some cases, the lack of the symmetric characteristics (e.g. S100A9, S100P) was observed suggesting that the two β^3^-amino acids are not interchangeable with each other, since not only the relative position of the side chains is important, but also the position of the preferred proteinogenic side chains related to the terminals.

### Investigating the interactions between the S100ome and the selected foldamers by fluorescence polarization

HTP (and also low-throughput) measurements generally need to be validated by an orthogonal approach to eliminate experimental artifacts [17]. We selected 11 foldamers (WL, IF, WW, YF, IL, VL, TW, RF, RR, TI, TM) based on the HTP HU assays, and after resynthesizing with a fluorescent label at the C-terminus (**Fig S13-15**, **Table S1**), the S100ome was tested against the labeled foldamers by direct fluorescence polarization (FP) (**Fig 3A**). In this assay, the association of the fluorescently labeled foldamer and the S100 protein of interest is monitored, through the change in polarization of the emitted light by the fluorophore upon the binding event. In direct FP, the presence of the fluorophore might change the binding affinity of some foldamers. While it would have been preferable to address the binding capacity of non-labelled foldamers by competitive FP [6], the limited solubility of the compounds and their low affinity did not allow us to set up a competitive assay. Nevertheless, we assumed that the fluorophore would affect all foldamers that target the same binding site to the same degree, because the foldamer scaffold is rigid. Defining the threshold of detection at the dissociation constant of 1 mM, we identified 87 interactions between the selected foldamer fragments and the S100ome out of 220 possible interactions (**Table 1, Fig S2-12**). The correlation between the two methodologies was quantitatively described by the calculated Pearson correlation coefficient (PCC = 0.36), resulting in a moderate correlation.

**Fig. 3.**
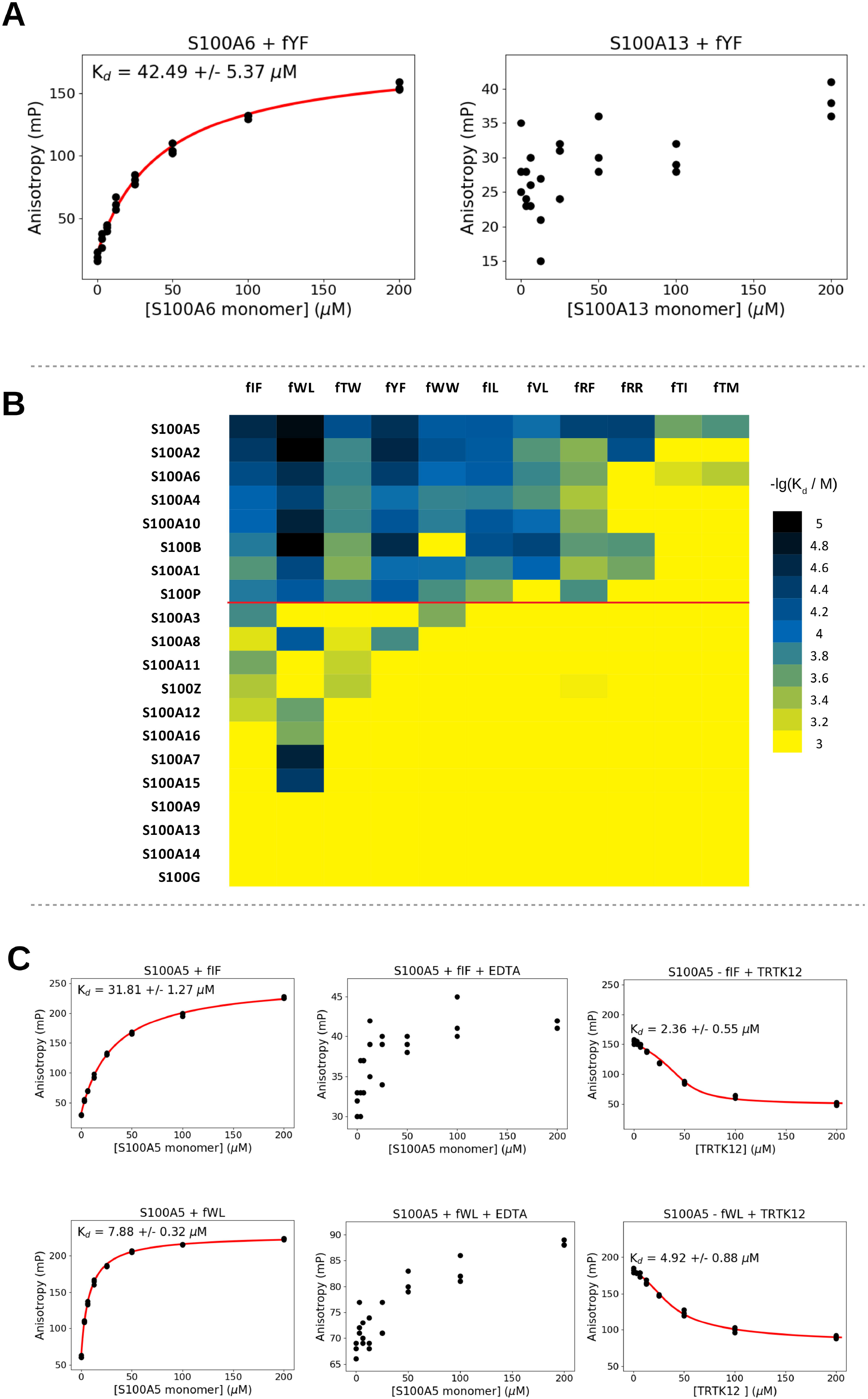
The interactions between the selected foldamers and S100 proteins measured by direct fluorescence polarization (FP) **Panel A:** The interactions between the S100ome and the labeled foldamer molecules were monitored by direct fluorescence polarization assay, in which the increase of the polarization (i.e. decrease of the rotation) caused by adding S100 proteins is indicative of the binding event, i.e. the association of the labeled foldamer – S100 complex is monitored. Dissociation constants were calculated by fitting the anisotropy values (mP) using quadratic equation with the ProFit program [6]. Left panel: S100A6 was added in various concentrations to the fluorescently labeled foldamer (fYF) and a significant binding event is observed. Right panel: S100A13 was added to the same foldamer and only a minor linear increase of the polarization was noticed confirming the results obtained by the HTP HU assays. **Panel B**: The -lg(K_d_) values of the interactions between the selected foldamers and the S100ome were depicted as a heat map. -lg(K_d_) ranges are color coded as shown on the right. The specificity-map of the S100ome towards the H14 library correlate well qualitatively with the results of the HTP HU measurements; i.e. S100 proteins (e.g. S100A2, S100A5, S100A6) interacting with numerous foldamer fragments in the HU assays exhibit the same behavior in direct FP measurements, meanwhile S100 members (e.g. S100G, S100A9, S100A13) imposing fewer interaction with the H14 library in the HU assays form weak, or no bound with the selected foldamers. It is noteworthy that based on the specificity map, the S100 proteins can be divided into two groups; one with numerous detected partners (upper part) and one with few or no detected partners (lower part). **Panel C:** Foldamer fragments bind to the hydrophobic binding pocket of S100 proteins in a calcium-dependent manner. Left panels: examples of direct titration of fIF and fWL in the presence of Ca^2+^ with S100A5, respectively, showing significant binding. Middle panels: the same titrations in the presence of EDTA resulted in the loss of binding event for both foldamer fragments, providing evidence that the S100-foldamer interactions are calcium-dependent. Right panels: Titrating the preformed S100 - labeled foldamer complex with an S100 binding peptide, TRTK12, competition between the labeled foldamer fragment and the unlabeled S100-binding peptide is observed in both cases, providing evidence that the foldamer fragments bind to the binding pocket of S100 proteins. K_d_ values were calculates as in Panel A using quadratic (left panels) and competitive binding equation (right panels), respectively.

**Table 1.** The dissociation constants given as mean ± SD of the selected foldamers and the S100ome measured by direct fluorescence polarization

|  | Kd (μM) |  |  |  |  |  |  |  |  |  |  |
| --- | --- | --- | --- | --- | --- | --- | --- | --- | --- | --- | --- |
|  | fIF | fRF | fTM | fTW | fWW | fTI | fIL | fRR | fVL | fWL | fYF |
| **S100A1** | 212 ± 23 | 413 ± 141 | >1000 | 329 ± 34 | 113 ± 33 | >1000 | 164 ± 41 | 278 ± 65 | 97 ± 21 | 53 ± 9 | 110 ± 17 |
| **S100A2** | 38 ± 2.6 | 339 ± 36 | >1000 | 173 ± 10 | 65 ± 2.0 | >1000 | 78 ± 15 | 61 ± 9.6 | 213 ± 183* | 11 ± 0.68 | 26 ± 3.7 |
| **S100A3** | 179 ± 14 | >1000 | >1000 | >1000 | 307 ± 37 | >1000 | >1000 | >1000 | >1000 | >1000 | >1000 |
| **S100A4** | 93 ± 4.1 | 462 ± 41 | >1000 | 182 ± 80 | 159 ± 17 | >1000 | 154 ± 8.3 | >1000 | 206 ± 15 | 46 ± 2.8 | 113 ± 9.0 |
| **S100A5** | 27 ± 0.91 | 46 ± 2.0 | 204 ± 16 | 62 ± 2.5 | 77 ± 4.9 | 270 ± 25 | 75 ± 6.2 | 45 ± 1.4 | 114 ± 9.3 | 14 ± 0.96 | 32 ± 1.7 |
| **S100A6** | 57 ± 3.2 | 285 ± 25 | 517 ± 99 | 161 ± 14 | 102 ± 5.5 | 699 ± 109 | 82 ± 9.3 | >1000 | 165 ± 22 | 30 ± 2.9 | 42 ± 5.7 |
| **S100A7** | >1000 | >1000 | >1000 | >1000 | >1000 | >1000 | >1000 | >1000 | >1000 | 23 ± 2.6 | >1000 |
| **S100A8** | 745 ± 79 | >1000 | >1000 | 769 ± 312 | >1000 | >1000 | >1000 | >1000 | >1000 | 74 ± 6 | 184 ± 73 |
| **S100A9** | >1000 | >1000 | >1000 | >1000 | >1000 | >1000 | >1000 | >1000 | >1000 | >1000 | >1000 |
| **S100A10** | 97 ± 4.3 | 320 ± 33 | >1000 | 170 ± 10 | 146 ± 9.0 | >1000 | 72 ± 2.8 | >1000 | 106 ± 9.4 | 23 ± 1.2 | 70 ± 6.0 |
| **S100A11** | 284 ± 26 | 998 ± 376 | >1000 | 574 ± 117 | >1000 | >1000 | >1000 | >1000 | >1000 | >1000 | >1000 |
| **S100A12** | 593 ± 68 | >1000 | >1000 | >1000 | >1000 | >1000 | >1000 | >1000 | >1000 | 254 ± 196* | >1000 |
| **S100A13** | >1000 | >1000 | >1000 | >1000 | >1000 | >1000 | >1000 | >1000 | >1000 | >1000 | >1000 |
| **S100A14** | >1000 | >1000 | >1000 | >1000 | >1000 | >1000 | >1000 | >1000 | >1000 | >1000 | >1000 |
| **S100A15** | >1000 | >1000 | >1000 | >1000 | >1000 | >1000 | >1000 | >1000 | >1000 | 39 ± 4.2 | >1000 |
| **S100A16** | >1000 | >1000 | >1000 | >1000 | >1000 | >1000 | >1000 | >1000 | >1000 | 304 ± 227* | >1000 |
| **S100B** | 138 ± 14 | 226 ± 10 | >1000 | 280 ± 29 | >1000 | >1000 | 63 ± 7.7 | 210 ± 45 | 50 ± 11 | 11 ± 1.3 | 28 ± 3.0 |
| **S100G** | >1000 | >1000 | >1000 | >1000 | >1000 | >1000 | >1000 | >1000 | >1000 | >1000 | >1000 |
| **S100P** | 136 ± 10 | 191 ± 29 | >1000 | 170 ± 17 | 190 ± 12 | >1000 | 330 ± 160 | >1000 | >1000 | 75 ± 10 | 79 ± 10 |
| **S100Z** | 483 ± 53 | 891 ± 199 | >1000 | 517 ± 102 | >1000 | >1000 | >1000 | >1000 | >1000 | >1000 | >1000 |

\*Due to the massive SD value compared to the mean, these $K_d$ values were not used in further calculations.

The affinity profile of the S100ome was depicted as a heat map (**Fig 3B**), using the K_d_ values determined in direct FP measurements fitted with a quadratic binding equation by the ProFit program [6]. Based on the affinities, the S100ome can be divided into two groups. The upper group shown in **Figure 3B** contains S100 proteins (S100A5, S100A2, S100A6, S100A4, S100A10, S100B, S100A1, S100P) with multiple detected interactions, which can be characterized by micromolar binding affinities. Meanwhile, the lower part of the heat map consists of members (S100A3, S100A14, S100A8, S100A11, S100Z, S100A13, S100A12, S100A16, S100A7, S100A15, S100G, S100A9) without a clear binding preference implying only a few or no partners amongst the selected foldamers.

To ensure that foldamer fragments bind to the hydrophobic binding groove of S100 proteins, which opens upon binding of calcium ions, we performed FP experiments on a selected S100 protein, S100A5, in the presence of EDTA and TRTK12, an S100-binding peptide of 12 amino acids. Our results showed that S100A5 is unable to interact with either fIF or fWL in the absence of Ca^2+^. Moreover, TRTK12 peptide competed with both foldamers with a K_d_ value similar to our previous results (**Fig 3C**) [6]. Based on the similarity and redundancy among the S100 family, it can be assumed that all S100 proteins bind the members of the H14 foldamer library through their hydrophobic binding pocket in a Ca^2+^-dependent manner.

### Mapping promiscuity in the S100ome

Assuming that the LSM library contains all the relevant binary combinations of amino acid side chains covering all the side chain preferences of S100 proteins, our data provide information about the binding promiscuity of each S100 member. Herein, we will refer to “promiscuity” as a parameter capturing both the broadness of exploration of the potential ligand space and the strength of binding to the recognized ligands. Based on the K_d_ values of the HU assays, we define a quantitative promiscuity term, that is calculated for each S100 protein by dividing its average association constant against the library by the strongest association constant among them (**eq. 1**).

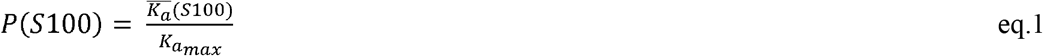

The determined promiscuity parameter represents the binding properties of each S100 family member against the applied foldamer library (**Fig 4A**). Higher values (e.g. in the case of S100A2 or S100A6) implicate a promiscuous behavior with numerous fragments to interact with, while lower values belong to S100 members (e.g. S100A7 or S100A13) with only a few, weak interactions or without a clear binding preference.

**Fig. 4.**
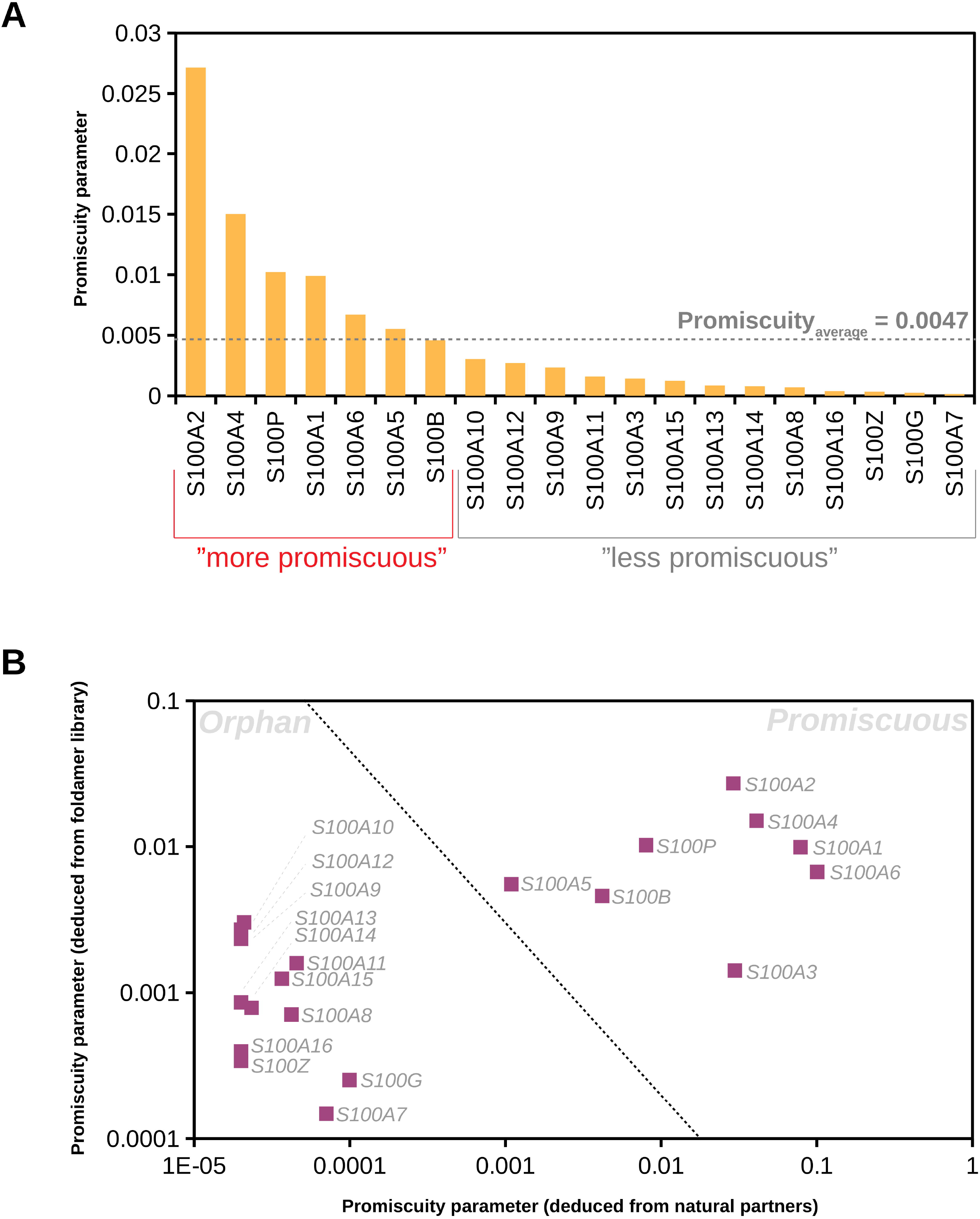
Promiscuity of the S100ome towards the H14 foldamer library. **Panel A:** Promiscuity values of the S100ome are defined towards the full H14 foldamer library (all the 256 possible combinations of 16 amino acids in two residues per foldamer building blocks) by summing up all the measured association constants and normalizing by the size of the library and the maximum of the measured association constants, and they are plotted on the y axis as a bar chart. The promiscuity average was arbitrarily chosen as threshold value for the promiscuous group (S100A2, S100A4, S100P, S100A1, S100A6, S100A5 and S100B), while the rest of the S100ome is less promiscuous exhibiting fewer binding events. **Panel B:** The Promiscuity values of the S100ome towards the foldamer library were plotted against the promiscuity values of the S100ome towards natural partners (based on 13 selected partners [6]) on logarithmic scale. It is shown that promiscuous (e.g. S100A6, S100B) and orphan (e.g. S100A13, S100G) members are clearly separated based on their promiscuity values, resulting in a good qualitative correlation between the two binding partner basis.

We also calculated the promiscuity parameter of each S100 member using our recently determined affinities of 13 model partners [6]. Despite we observed a moderate correlation between the promiscuities based on the artificially selected small number of natural partners and the foldamer library (PCC = 0.45), the two groups of promiscuous and orphan members could be clearly separated (**Fig 4B**).

## Discussion

### High-throughput holdup screening with foldamer libraries is a potent tool for specificity profiling of protein families

Using the H14 LSM library, the chemical-binding preferences of the S100ome were screened effectively by the application of a HTP-HU assay, in which numerous strongly interacting foldamers were identified. When considering our overall results, the quantity and quality of the selected foldamer residues was utilized to create the specificity map of the overall S100ome. The detected enrichment of the highly hydrophobic and/or aromatic residues on the interacting surface is not a unique feature of the foldamers; moreover, their side chain binding propensities are biomimetic. Certain aromatic and aliphatic amino acids (i.e. Trp/Phe/Tyr and Leu/Ile/Val) were especially favored on the binding interface and these findings are in line with literature data from protein-protein interaction interface databases [18]. In general, selective recognition of ligands can be explained with the unique binding patterns of the protein interfaces; therefore, the side chain frequency levels can be different even for proteins having considerably similar structures. Importantly, as other foldamer libraries with different constitution (i.e. the constitutional and/or spatial conformation of the β^3^-amino acid side chains in the foldamer fragments) are available (e.g. the H12 foldamer library), the affinity of the individual S100 members towards the foldamer libraries can vary. Therefore, it would be interesting to screen the S100ome against other foldamer libraries, which could reveal additional relationships between the S100 members through their binding properties, providing a more refined specificity map of the family.

To validate the detected interactions, an orthogonal biophysical method, direct FP technique was used. Importantly, the affinity profile of the S100ome against the selected foldamers shows good correlation with the specificity map of the S100ome from our previous work using natural S100 partners [6]. While S100 proteins with multiple natural interaction partners (e.g. S100B, S100A6) are keen on binding foldamers, S100 proteins without a clear binding preference (S100G, S100A13) can barely interact with LSMs, either because they only bind to proteins or peptides presenting a different conformation, or because they do not naturally bind to proteins.

Importantly, as S100 proteins are potential therapeutic targets, the concatenation of the smaller foldamer fragments screened here by the HTP HU assay might actually lead to highly specific and strong ligands, paving the way to rational drug design.

### The promiscuity of the full S100ome is explained using the foldamer library

Promiscuity (or its complementary notion, specificity) within a protein family can hardly be defined against natural partners, owing to still potentially unknown interactions. However, using the LSM foldamer library against the S100ome to screen the binding properties within the protein family, promiscuity can be defined for each member against the actual library, which eventually may approximate the real, yet undefined promiscuity profile. The promiscuity parameter values defined in this study for each S100 member against the foldamer library are in good correlation with previous works [6, 13, 19–25]. Promiscuous S100 proteins with several known cellular partners (e.g. S100A6 or S100A4) show more interactions towards the members of the foldamer library, thus displaying a higher value of their promiscuity parameter. Orphan S100 proteins without a clear intra- or extracellular binding preference (e.g. S100A16 or S100Z) exhibit less interactions with lower binding affinity, which is represented by a lower value of the promiscuity parameter.

Orphan members do not interact considerably with the members of the H14 foldamer library, thus suggesting that these S100 proteins might lack the ability to interact with proteins in the real cellular environment and rather play a role in the Ca^2+^-homeostasis [26]. It is still possible that the less promiscuous S100 proteins have highly specific, yet undiscovered natural interaction partners that may adopt a drastically different, non-helical conformation, explaining their lack of preference for the H14 helical foldamer fragments.

While, in principle, functional redundancy within the S100ome can only be interpreted with natural partners, screening the S100ome against ‘non-natural’ libraries constitutes a powerful approach to draw a more detailed and refined picture about binding properties within the family. The promiscuity of the S100 proteins observed herein against the foldamer library, may have high relevance for their actual interactome in the real cellular environment.

## Materials and Methods

### S100 protein expression and purification

S100 proteins (UniProt accession codes: S100A1: P23297, S100A2: P29034, S100A3: P33764, S100A4: P26447, S100A5: P33763, S100A6: P06703, S100A7: P31151, S100A8: P05109, S100A9: P06702, S100A10: P60903, S100A11: P31949, S100A12: P80511, S100A13: Q99584, S100A14: Q9HCY8, S100A15: Q86SG5, S100A16: Q96FQ6, S100B: P04271, S100G: P29377, S100P: P25815 and S100Z: Q8WXG8) were expressed and purified with N-terminal His_6_-tag as described previously [27]. Briefly, S100 proteins were cloned into a modified pET15b vector with a TEV protease cleavable N-terminal His_6_-tag and expressed in *Escherichia* coli BL21 (DE3) cells, followed by Ni^2+^-affinity chromatography. For HU assay, S100 proteins were further purified by either hydrophobic interaction chromatography or ion exchange chromatography without the cleavage of the N-terminal His_6_-tag applying standard conditions [27]. For direct FP measurements, the N-terminal His_6_-tag was cleaved, and the S100 proteins were purified by hydrophobic interaction chromatography, ion exchange chromatography or size exclusion chromatography [27]. The quality of the recombinant proteins was checked by SDS-PAGE analysis in all cases. The concentration of the recombinant S100 proteins was determined by UV spectrophotometry using the absorbance of Tyr and Trp residues.

### Synthesis and purification of the foldamer libraries

The foldamer libraries were synthetized and purified as described previously [28]. Briefly, the 256-memberd library was divided to four sublibraries (aromatic, charged, apolar, non-charged polar) containing 64 members. The libraries were synthetized with a CEM liberty 1 microwave peptide synthesizer using HATU (1-[bis(dimethylamino)methylene]-1H-1,2,3-triazolo[4,5-b]-pyridinium-3-oxid hexafluorophosphate) as coupling agent following Fmoc strategy by coupling aminocyclohexanecarboxylic acids and β^3^-amino acids. After cleavage of the sublibraries, the samples were lyophilized and the mixtures of foldamers were purified by RP-HPLC (Phenomenex Luna C18, 250 × 10 mm), followed by HPLC-MS identification. The purity and equimolarity of the foldamer libraries were checked by HPLC-MS.

### Synthesis and purification of labeled foldamer sequences

Individual foldamers were synthetized manually using solid-phase peptide synthesis with Fmoc strategy applying HATU as coupling agent [12]. Coupling of the 5(6)-carboxyfluorescein to the ε-amino group of a Lys attached to the C-terminus of the foldamers was carried out as the last step of the synthesis. The crude foldamers were cleaved from the resin and then, the samples were precipitated in diethyl ether and purified by RP-HPLC (Phenomenex Jupiter C18, 250 × 10 mm). Purity was confirmed by HPLC-MS. The concentration of the foldamers was determined by UV-spectrophotometry using the absorbance of 5(6)-carboxyfluorescein.

### Synthesis and purification of the peptide TRTK12

The TRTK12 peptide was synthetized as described previously [6]. Briefly, the peptide was chemically synthetized by solid phase peptide synthesis with a PS3 peptide synthesizer (Protein technologies, Tucson, AZ, USA) using Fmoc/tBu strategy, and purified by RP-HPLC using a Jupiter 300 Å C_18_ column.

### Holdup assay

Screening the interaction between the foldamer libraries and the S100ome was performed by holdup assays as described previously [28]. Briefly, S100 proteins were immobilized in a buffer containing 20 mM HEPES pH 7.5, 150 mM NaCl, 2 mM CaCl_2_, 1 mM TCEP on Co^2+^-affinity resin (~2 mg protein / ml resin concentration) via the N-terminal His_6_-tag followed by the addition of the foldamer libraries. After incubation, the resin was centrifuged (*Pierce™ Spin Cups – paper filter, Thermo Fisher Scientific*) to separate the unbound fraction of the library. S100 – foldamer complexes were eluted with 250 mM imidazole added to the binding buffer. Negative controls were prepared using the procedure described above in the absence of the His_6_-tagged protein. The flow-through and eluted fractions were analyzed by HPLC-MS. Quantitative evaluation of the HPLC-MS chromatograms were performed with Thermo Xcalibur software. Bound fractions (F_B_) were calculated by the following equation (**eq. 2**) from the loss of intensity of the foldamer fragments (AUC_protein_) in the flow-through fractions compared to the control samples (AUC_control_).

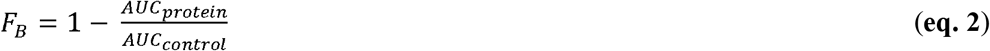

K_d_ values of the i^th^ foldamer were calculated with the formula described below (**eq. 3**) using the calculated bound fractions and assuming that S100 proteins were quantitatively immobilized on the resin in 64 μM.

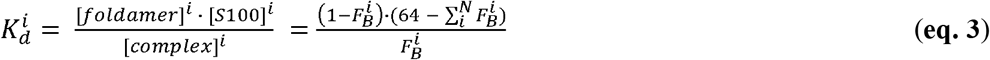

AUC value of the foldamer fragment in the eluted fraction was determined for the purpose of qualitative comparison, but was not used for further calculations.

### Calculation of amino acid preference

For each 16 amino acid, a summarized F_B_ (F_B_^aa^) was calculated by the following equation in the instances of all S100 porteins:

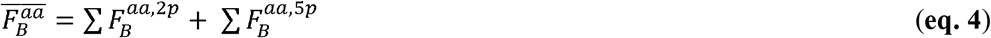

In which F_B_^aa,2p^ and F_B_^aa,5p^ are fraction bound values of foldamer fragments containing the proteogenic sidechain of interest in the 2^nd^ or 5^th^ position, respectively. The amino acids were further categorized into five groups (aromatic: F, W, Y; aliphatic: A, I, L, M, V; polar: N, Q, S, T; acidic: D, E; basic: K, R), and the root fraction bound values (F_B_^root^) were calculated for each group in the case of all S100 proteins by the following equation:

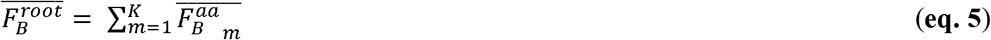

In which K is the number of amino acids in the individual groups.

### Fluorescence polarization assay

In direct fluorescence polarization assays, S100 proteins were diluted in a buffer containing 50 nM labeled foldamer, 20 mM HEPES pH 7.5, 150 mM NaCl, 1 mM CaCl_2_, 0.5 mM TCEP and 0.01% Tween20. The dilution series (50 μl) were divided into three technical repeats and transferred (15 μl) to a 384-well microplate. In competitive fluorescence polarization assays, the buffer applied in direct measurements was supplemented with the S100 protein of interest to reach a saturation of 60-80%. This mixture was titrated with the competitor (i.e. the unlabeled peptide). Fluorescence polarization was measured in 8 different S100 concentrations (one of which contained no S100 protein) on a Synergy H4 plate reader using 485 ± 20 nm and 528 ± 20 nm band-pass filters for excitation and emission, respectively. The K_d_ values were obtained by fitting the data from the FP measurements with the python-based ProFit software using quadratic and competitive binding equation for direct and competitive FP, respectively [6]. The detection threshold was based on two parameters. First, we rejected all fitted dissociation constants above 1 mM. Second, we also rejected all fitted data where the experimental window was significantly lower (< 80 mP) or higher (> 350 mP), compared to other, stronger interactions of the same labeled foldamer.

### Correlation between holdup and FP

The correlation between the holdup assay and the fluorescence polarization was quantitatively described by the Pearson correlation coefficient (PCC) using the standard formula.

### Calculation of promiscuity

Promiscuity, defined as a number between 0 (no interaction with any member) and 1 (the strongest interaction with all the members in the library) was calculated for each S100 protein according to **eq. 1** by averaging the measured apparent dissociation constants which was then normalized to the strongest K_d_ value measured against the foldamer library. For calculating the promiscuity of S100 members, based on natural partners, we used our previously measured interactomic map and we used a kNN approach, based on the average of the 2 nearest neighbor in the UPGMA clustering to fill the few missing binding parameters [6]. The Pearson correlation coefficient (PCC) was calculated between the two promiscuities using the standard formula.

#### Accession numbers

**S100A1**: UniProt accession number **<u>P23297</u>**, **S100A2**: UniProt accession number **<u>P29034</u>**, **S100A3**: UniProt accession number **<u>P33764</u>**, **S100A4**: UniProt accession number **<u>P26447</u>**, **S100A5**: UniProt accession number **<u>P33763</u>**, **S100A6**: UniProt accession number **<u>P06703</u>**, **S100A7**: UniProt accession number **<u>P31151</u>**, **S100A8**: UniProt accession number **<u>P05109</u>**, **S100A9**: UniProt accession number **<u>P06702</u>**, **S100A10**: UniProt accession number **<u>P60903</u>**, **S100A11**: UniProt accession number **<u>P31949</u>**, **S100A12**: UniProt accession number **<u>P80511</u>**, **S100A13**: UniProt accession number **<u>Q99584</u>**, **S100A14**: UniProt accession number **<u>Q9HCY8</u>**, **S100A15**: UniProt accession number **<u>Q86SG5</u>**, **S100A16**: UniProt accession number **<u>Q96FQ6</u>**, **S100B**: UniProt accession number **<u>P04271</u>**, **S100G**: UniProt accession number **<u>P29377</u>**, **S100P**: UniProt accession number **<u>P25815</u>**, **S100Z**: UniProt accession number **<u>Q8WXG8</u>**.

## List of abbreviations

AUC: area under curve
FP: fluorescence polarization
HTP: high-throughput
LC-MS: liquid chromatography - mass spectrometry
HU: holdup
PPI: protein-protein interactions
RP-HPLC: reverse phased high performance liquid chromatography
SDS-PAGE: sodium dodecyl sulfate polyacrylamide gel electrophoresis

## Acknowledgement

This work was supported by the National Research Development and Innovation Fund of Hungary (K119359 to LN). MAS was supported through the New National Excellence Program of the Hungarian Ministry of Human Capacities (UNKP-18-2). We also acknowledge the FIEK_16-1-2016-0005 and VEKOP-2.3.3-15-2016-00011 grants. This work was completed as part of the ELTE Thematic Excellence Programme 2020 supported by the National Research, Development and Innovation Office (TKP2020-IKA-05).

## Author contribution

MAS carried out protein expression, FP experiments, FP data analysis, calculations, and wrote the paper. ÉB prepared the foldamers, executed the HU assays, analyzed the HU data and wrote the paper. BM, LR and EB contributed by carrying out foldamer synthesis and protein expression, HU experiments and data analysis. GG analyzed the data and wrote the manuscript. GT, TAM and LN supervised the research, analyzed the data and wrote the paper.

## Supplementary Figure and Table Legends

**Fig S1**. The binding affinities of the S100ome towards the H14 foldamer library measured by a high-throughput holdup assay. The calculated dissociation constants were depicted as a heat map on a linear scale. K_d_ ranges are color-coded as shown on the right. The missing S100 proteins can be found in following reference (Tököli et al., 2020).

**Fig S2.** The interactions between the S100ome and fIF as measured by FP. Dissociation constants were calculated by fitting the anisotropy values (mP) using quadratic equation with the ProFit program.

**Fig S3.** The interactions between the S100ome and fIL as measured by FP. Dissociation constants were calculated by fitting the anisotropy values (mP) using quadratic equation with the ProFit program.

**Fig S4.** The interactions between the S100ome and fRF as measured by FP. Dissociation constants were calculated by fitting the anisotropy values (mP) using quadratic equation with the ProFit program.

**Fig S5.** The interactions between the S100ome and fRR as measured by FP. Dissociation constants were calculated by fitting the anisotropy values (mP) using quadratic equation with the ProFit program.

**Fig S6**. The interactions between the S100ome and fTI as measured by FP. Dissociation constants were calculated by fitting the anisotropy values (mP) using quadratic equation with the ProFit program.

**Fig S7**. The interactions between the S100ome and fTM as measured by FP. Dissociation constants were calculated by fitting the anisotropy values (mP) using quadratic equation with the ProFit program.

**Fig S8**. The interactions between the S100ome and fTW as measured by FP. Dissociation constants were calculated by fitting the anisotropy values (mP) using quadratic equation with the ProFit program.

**Fig S9**. The interactions between the S100ome and fVL as measured by FP. Dissociation constants were calculated by fitting the anisotropy values (mP) using quadratic equation with the ProFit program.

**Fig S10**. The interactions between the S100ome and fWL as measured by FP. Dissociation constants were calculated by fitting the anisotropy values (mP) using quadratic equation with the ProFit program.

**Fig S11**. The interactions between the S100ome and fWW as measured by FP. Dissociation constants were calculated by fitting the anisotropy values (mP) using quadratic equation with the ProFit program.

**Fig S12**. The interactions between the S100ome and fYF as measured by FP. Dissociation constants were calculated by fitting the anisotropy values (mP) using quadratic equation with the ProFit program.

**Fig S13**. The structures of the selected foldamer sequences labeled with 5(6)-carboxyfluorescein. Foldamers were coupled to the fluorescence dye through two glycine residues. R1 and R2 represent proteogenic side chains at second and fifth positions, respectively.

**Fig S14**. The MS spectra and the HPLC chromatogram of fWW (Column: Phenomenex Luna C18 (250 × 4.6 mm, particle size: 5 micron, pore size: 100Å); Gradient: 5-80% 20min 1.2 mL min-1); fWL (Column: Phenomenex Luna C18 (250 × 4.6 mm, particle size: 5 micron, pore size: 100Å); Gradient: 60-80% 20min 1.2 mL min-1); fYF (Column: Phenomenex Luna C18 (250 × 4.6 mm, particle size: 5 micron, pore size: 100Å); Gradient: 5-80% 20min 1.2 mL min-1); fIF (Column: Phenomenex Luna C18 (250 × 4.6 mm, particle size: 5 micron, pore size: 100Å); Gradient: 40-60% 20min 1.2 mL min-1); fTW (Column: Phenomenex Luna C18 (250 × 4.6 mm, particle size: 5 micron, pore size: 100Å); Gradient: 5-80% 20min 1.2 mL min-1) and fRF (Column: Phenomenex Luna C18 (250 × 4.6 mm, particle size: 5 micron, pore size: 100Å); Gradient: 5-80% 20min 1.2 mL min-1).

**Fig S15**. The MS spectra and the HPLC chromatogram of fII (Column: Phenomenex Luna C18 (250 × 4.6 mm, particle size: 5 micron, pore size: 100Å); Gradient: 40-60% 20min 1.2 mL min-1); fVL (Column: Phenomenex Luna C18 (250 × 4.6 mm, particle size: 5 micron, pore size: 100Å); Gradient: 40-70% 30min 1.2 mL min-1); fRR (Column: Phenomenex Luna C18 (250 × 4.6 mm, particle size: 5 micron, pore size: 100Å); Gradient: 5-80% 20min 1.2 mL min-1); fTI (Column: Phenomenex Luna C18 (250 × 4.6 mm, particle size: 5 micron, pore size: 100Å); Gradient: 5-80% 30min 1.2 mL min-1); fTM (Column: Phenomenex Luna C18 (250 × 4.6 mm, particle size: 5 micron, pore size: 100Å); Gradient: 30-50% 20min 1.2 mL min-1) and fIL (Column: Phenomenex Luna C18 (250 × 4.6 mm, particle size: 5 micron, pore size: 100Å); Gradient: 40-70% 30min 1.2 mL min-1).

**Table S1**. Summary of the ESI-MS data for the selected foldamers labeled with the fluorescence dye.

## References

[1] Donato R. S100: a multigenic family of calcium-modulated proteins of the EF-hand type with intracellular and extracellular functional roles. Int J Biochem Cell Biol. 2001;33:637–68.

[2] Donato R. Functional roles of S100 proteins, calcium-binding proteins of the EF-hand type. Biochim Biophys Acta. 1999;1450:191–231.

[3] Bresnick AR, Weber DJ, Zimmer DB. S100 proteins in cancer. Nat Rev Cancer. 2015;15:96–109.

[4] Chen H, Xu C, Jin Qe, Liu Z. S100 protein family in human cancer. American journal of cancer research. 2014;4:89–115.

[5] Bresnick AR. S100 proteins as therapeutic targets. Biophysical Reviews. 2018;10:1617–29.

[6] Simon MA, Ecsedi P, Kovacs GM, Poti AL, Remenyi A, Kardos J, et al. High-throughput competitive fluorescence polarization assay reveals functional redundancy in the S100 protein family. FEBS J. 2020;287:2834–46.

[7] Pelay-Gimeno M, Glas A, Koch O, Grossmann TN. Structure-Based Design of Inhibitors of Protein-Protein Interactions: Mimicking Peptide Binding Epitopes. Angew Chem Int Ed Engl. 2015;54:8896–927.

[8] Cabrele C, Martinek TA, Reiser O, Berlicki Ł. Peptides Containing β-Amino Acid Patterns: Challenges and Successes in Medicinal Chemistry. Journal of Medicinal Chemistry. 2014;57:9718–39.

[9] Checco JW, Gellman SH. Targeting recognition surfaces on natural proteins with peptidic foldamers. Curr Opin Struct Biol. 2016;39:96–105.

[10] Guichard G, Huc I. Synthetic foldamers. Chemical Communications. 2011;47:5933–41.

[11] Tököli A, Mag B, Bartus É, Wéber E, Szakonyi G, Simon MA, et al. Proteomimetic surface fragments distinguish targets by function. Chemical Science. 2020;11:10390–8.

[12] Bartus E, Hegedus Z, Weber E, Csipak B, Szakonyi G, Martinek TA. De Novo Modular Development of a Foldameric Protein-Protein Interaction Inhibitor for Separate Hot Spots: A Dynamic Covalent Assembly Approach. ChemistryOpen. 2017;6:236–41.

[13] Wheeler LC, Anderson JA, Morrison AJ, Wong CE, Harms MJ. Conservation of Specificity in Two Low-Specificity Proteins. Biochemistry. 2018;57:684–95.

[14] Gogl G, Biri-Kovacs B, Durbesson F, Jane P, Nomine Y, Kostmann C, et al. Rewiring of RSK-PDZ Interactome by Linear Motif Phosphorylation. J Mol Biol. 2019;431:1234–49.

[15] Vincentelli R, Luck K, Poirson J, Polanowska J, Abdat J, Blemont M, et al. Quantifying domain-ligand affinities and specificities by high-throughput holdup assay. Nat Methods. 2015;12:787–93.

[16] Luck K, Trave G. Phage display can select over-hydrophobic sequences that may impair prediction of natural domain-peptide interactions. Bioinformatics. 2011;27:899–902.

[17] Finak G, Gottardo R. Promises and Pitfalls of High-Throughput Biological Assays. In: Carugo O, Eisenhaber F, editors. Data Mining Techniques for the Life Sciences. New York, NY: Springer New York; 2016. p. 225–43.

[18] Watkins AM, Bonneau R, Arora PS. Side-Chain Conformational Preferences Govern Protein– Protein Interactions. Journal of the American Chemical Society. 2016;138:10386–9.

[19] Biri-Kovács B, Kiss B, Vadászi H, Gógl G, Pálfy G, Török G, et al. Ezrin interacts with S100A4 via both its N- and C-terminal domains. PLoS ONE. 2017;12:e0177489–e.

[20] Ecsédi P, Billington N, Pálfy G, Gógl G, Kiss B, Bulyáki É, et al. Multiple S100 protein isoforms and C-terminal phosphorylation contribute to the paralog-selective regulation of nonmuscle myosin 2 filaments. Journal of Biological Chemistry. 2018;293:14850–67.

[21] Ecsédi P, Kiss B, Gógl G, Radnai L, Buday L, Koprivanacz K, et al. Regulation of the Equilibrium between Closed and Open Conformations of Annexin A2 by N-Terminal Phosphorylation and S100A4-Binding. Structure. 2017;25:1195–207.

[22] Fernandez-Fernandez MR, Rutherford TJ, Fersht AR. Members of the S100 family bind p53 in two distinct ways. Protein Science. 2008;17:1663–70.

[23] Gógl G, Alexa A, Kiss B, Katona G, Kovács M, Bodor A, et al. Structural Basis of Ribosomal S6 Kinase 1 (RSK1) Inhibition by S100B Protein. Journal of Biological Chemistry. 2015;291:11–27.

[24] Liu Y, Myrvang HK, Dekker LV. Annexin A2 complexes with S100 proteins: Structure, function and pharmacological manipulation. British Journal of Pharmacology. 2015;172:1664–76.

[25] Shimamoto S, Kubota Y, Yamaguchi F, Tokumitsu H, Kobayashi R. Ca 2+ /S100 proteins act as upstream regulators of the chaperone-associated ubiquitin ligase chip (c terminus of hsc70-interacting protein). Journal of Biological Chemistry. 2013;288:7158–68.

[26] Schwaller B. Cytosolic Ca2+ Buffers. Cold Spring Harbor Perspectives in Biology. 2010;2:a004051–a.

[27] Kiss B, Ecsédi P, Simon M, Nyitray L. Isolation and Characterization of S100 Protein-Protein Complexes. Methods in Molecular Biology. 2019;1929:325–38.

[28] Bartus É, Olajos G, Schuster I, Bozsó Z, Deli MA, Veszelka S, et al. Structural Optimization of Foldamer-Dendrimer Conjugates as Multivalent Agents against the Toxic Effects of Amyloid Beta Oligomers. Molecules. 2018;23:2523.

[29] Kiss B, Duelli A, Radnai L, Kekesi KA, Katona G, Nyitray L. Crystal structure of the S100A4-nonmuscle myosin IIA tail fragment complex reveals an asymmetric target binding mechanism. Proc Natl Acad Sci U S A. 2012;109:6048–53.

