## Supplementary figures and images for "Binding Profile Mapping of the S100 Protein Family Using a High-throughput Local Surface Mimetic Holdup Assay"

### Supplemental Fig 1

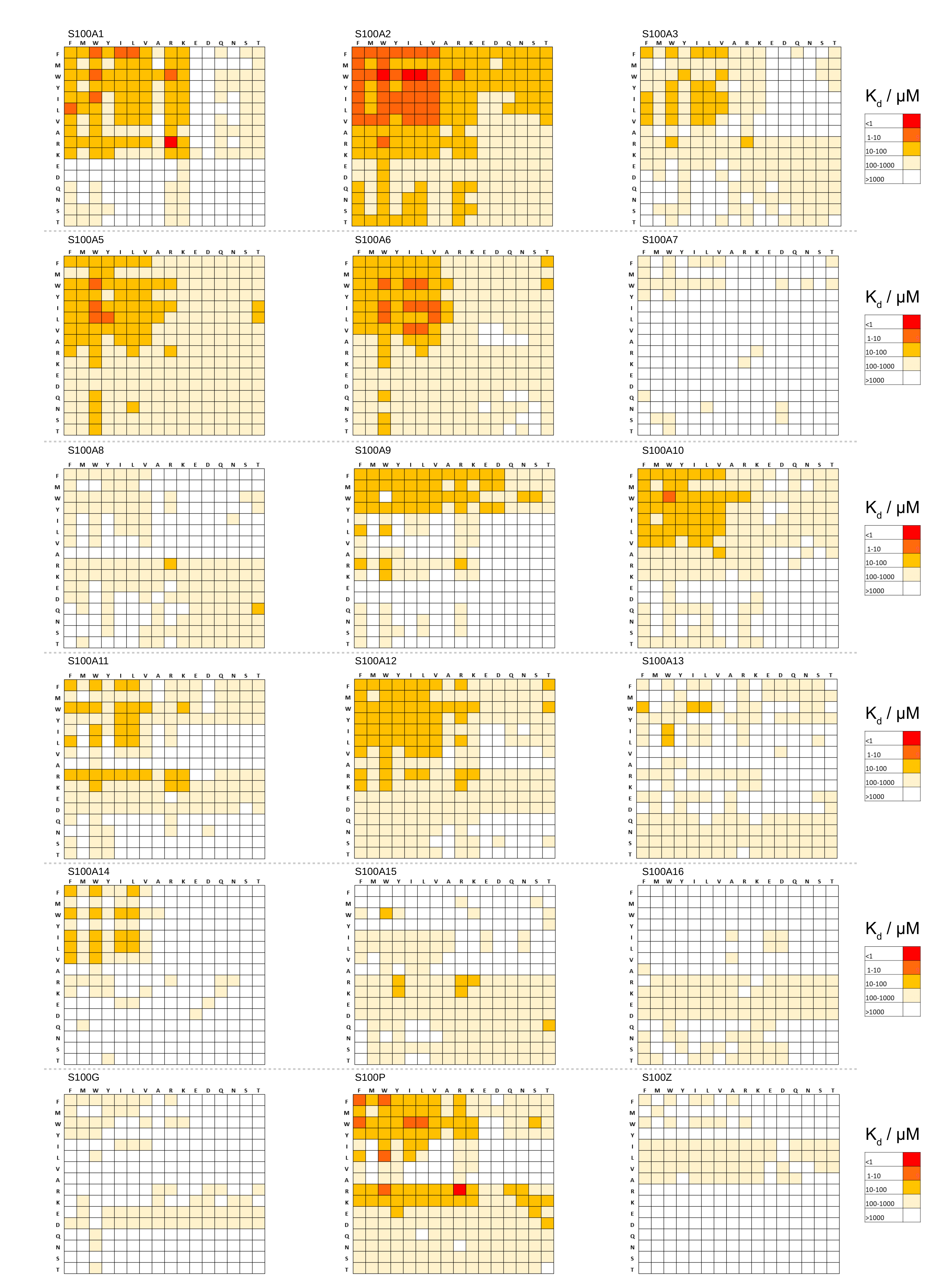

### Supplemental Fig 2

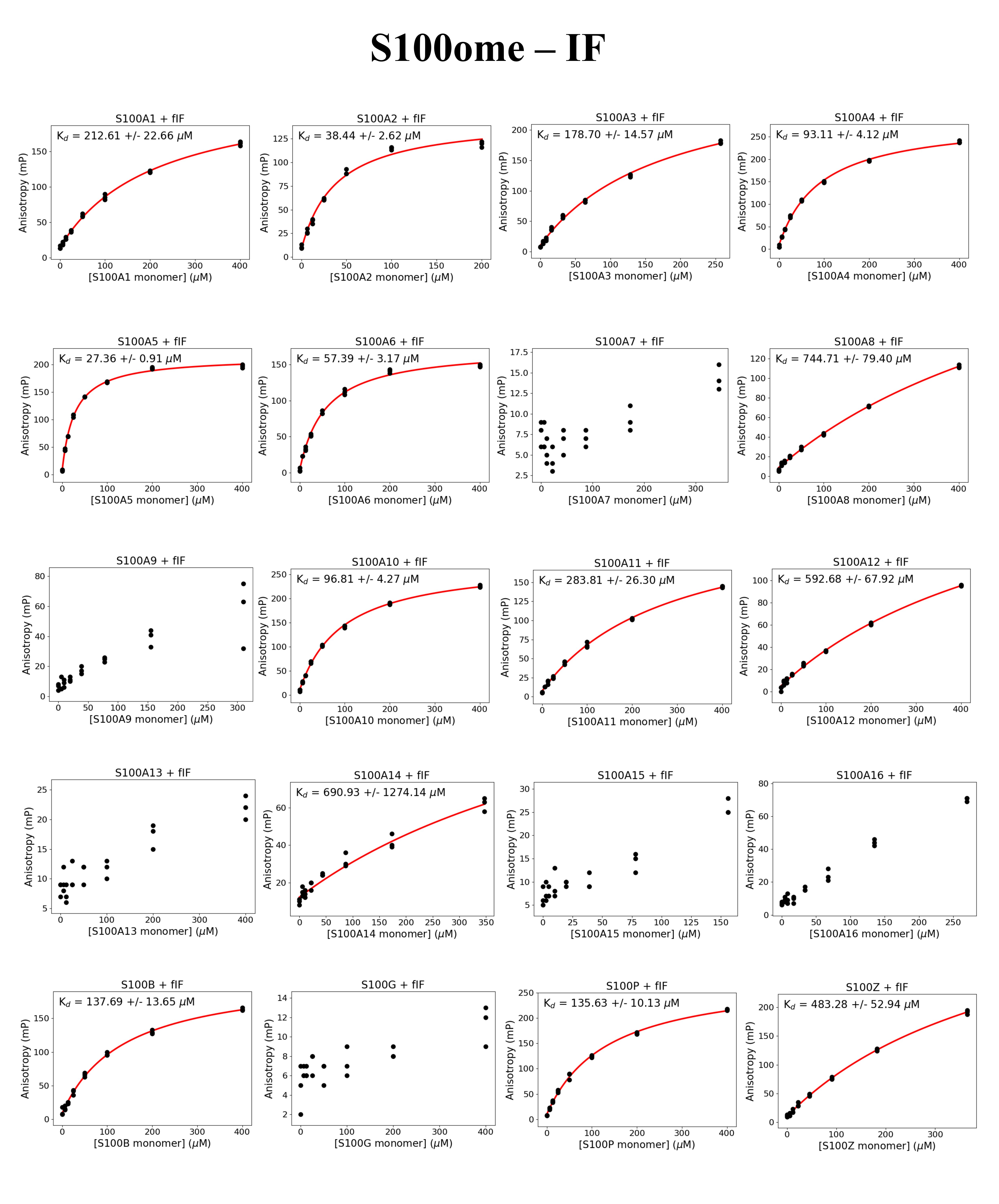

### Supplemental Fig 3

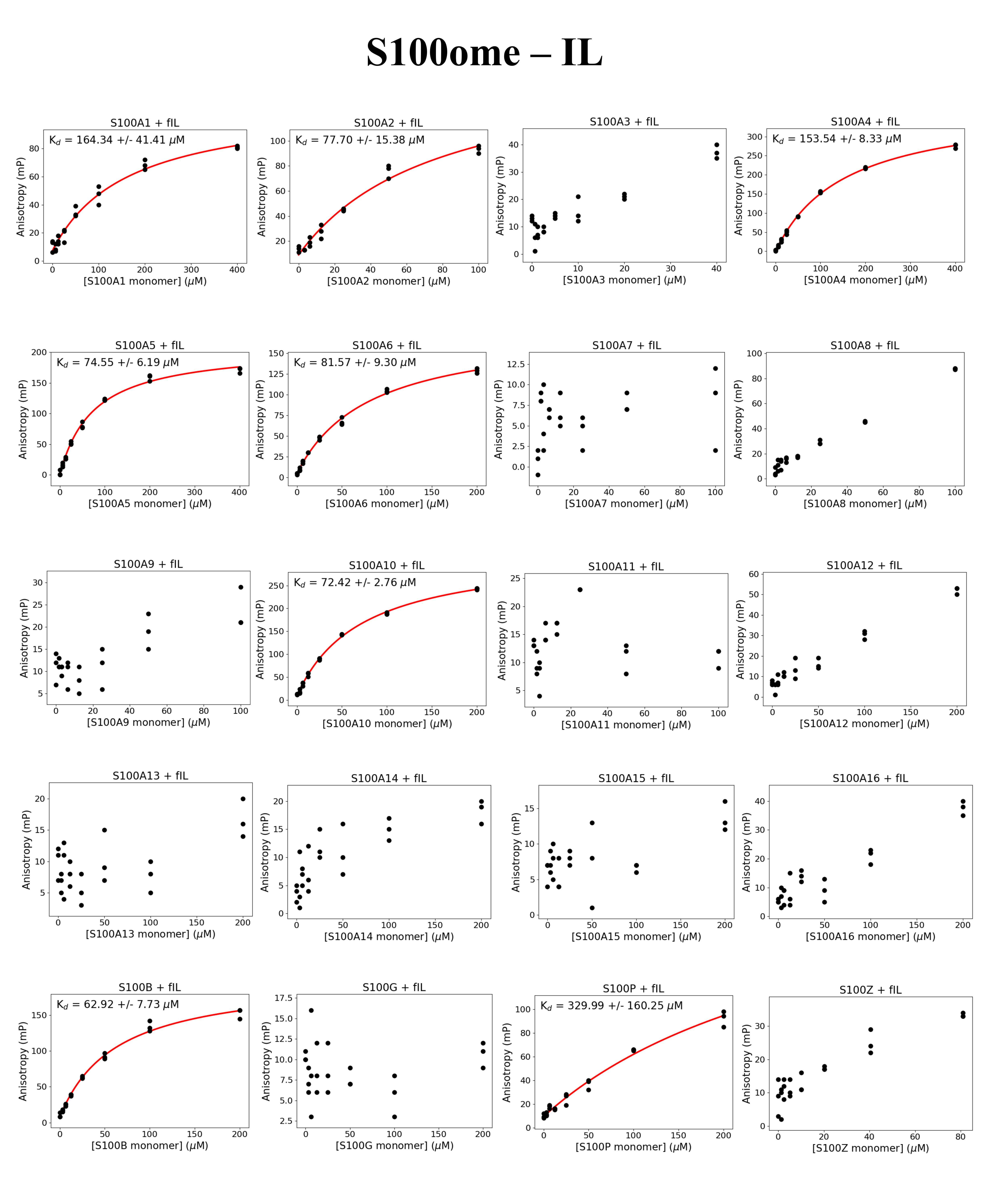

### Supplemental Fig 4

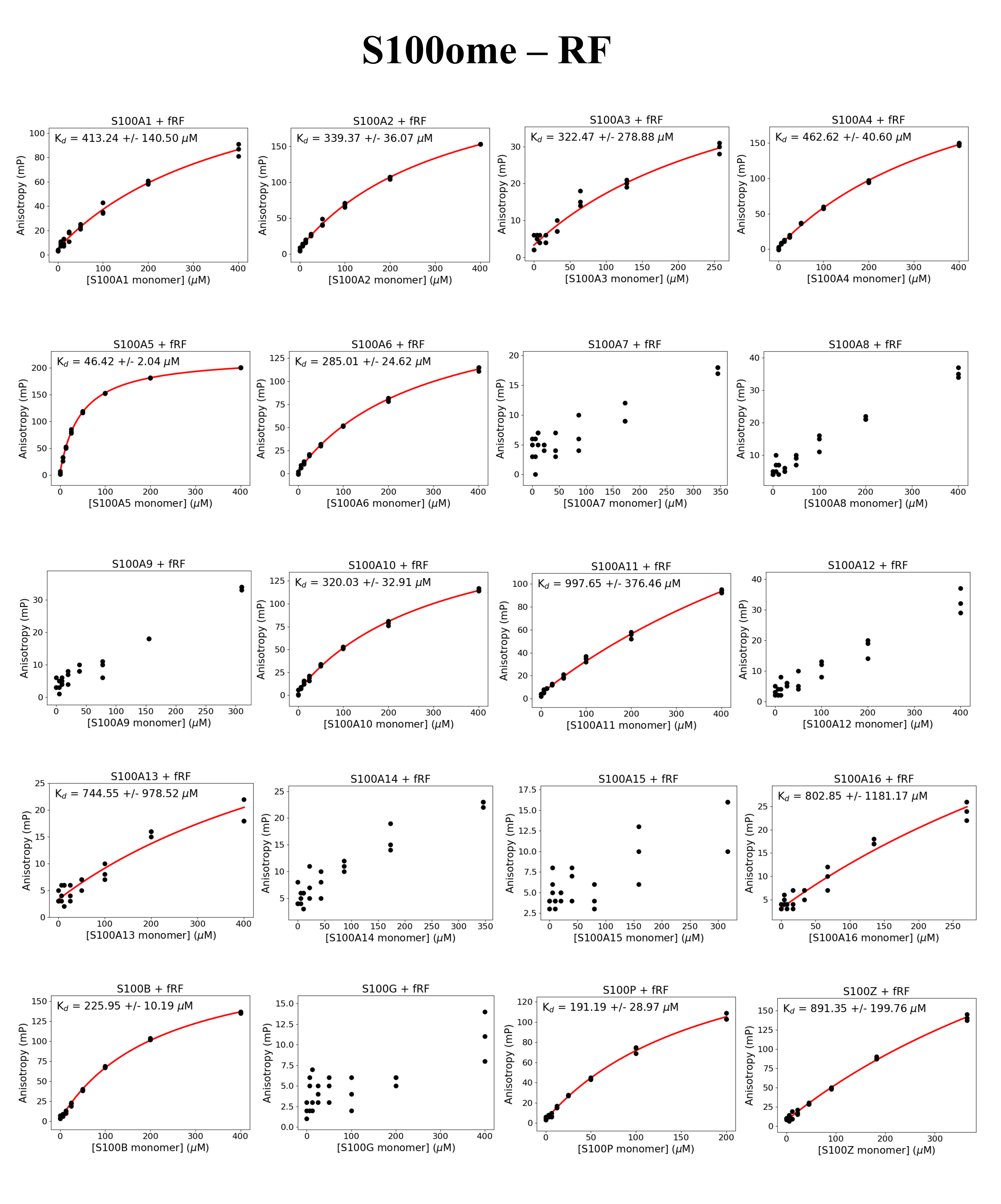

### Supplemental Fig 5

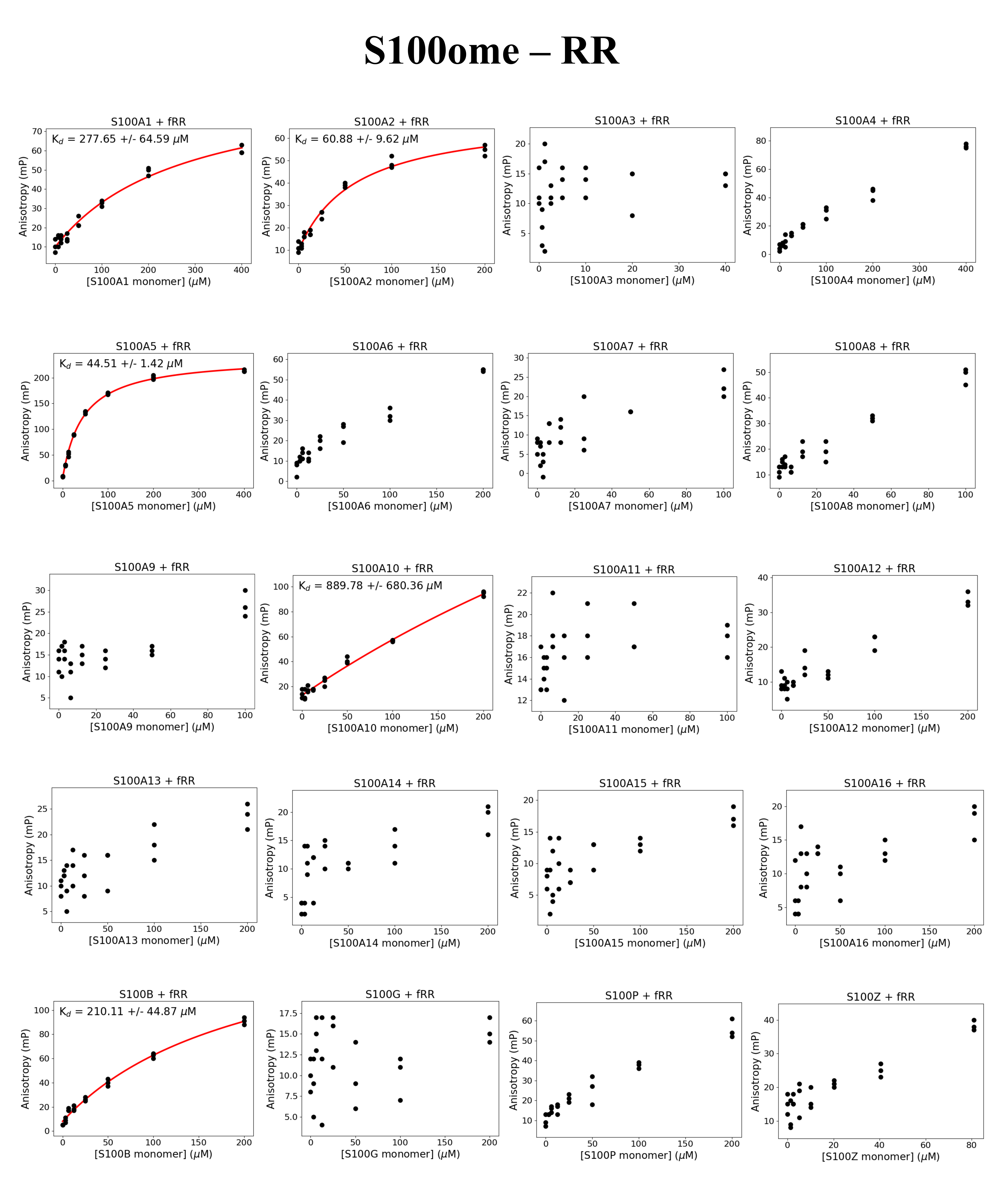

### Supplemental Fig 6

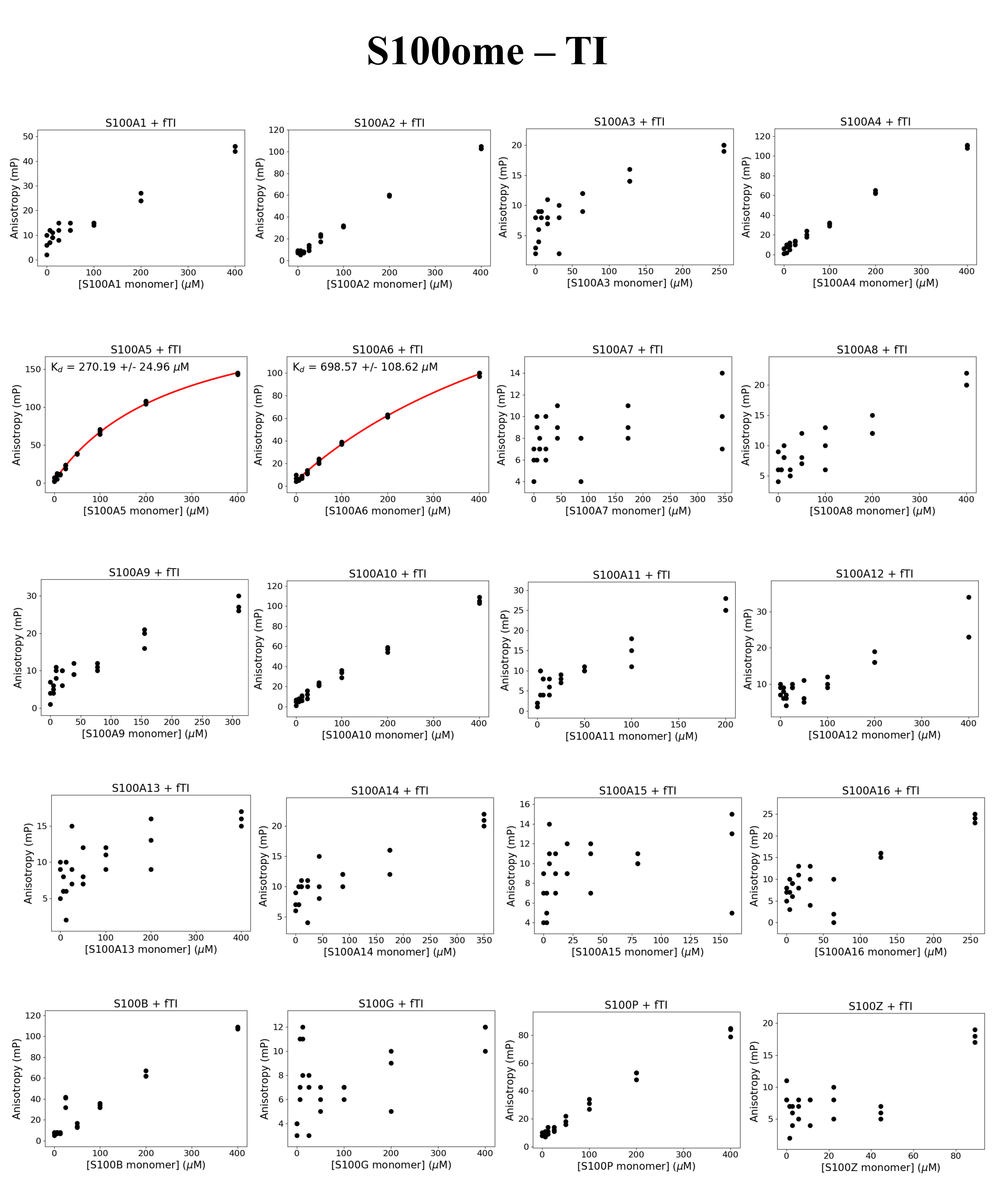

### Supplemental Fig 7

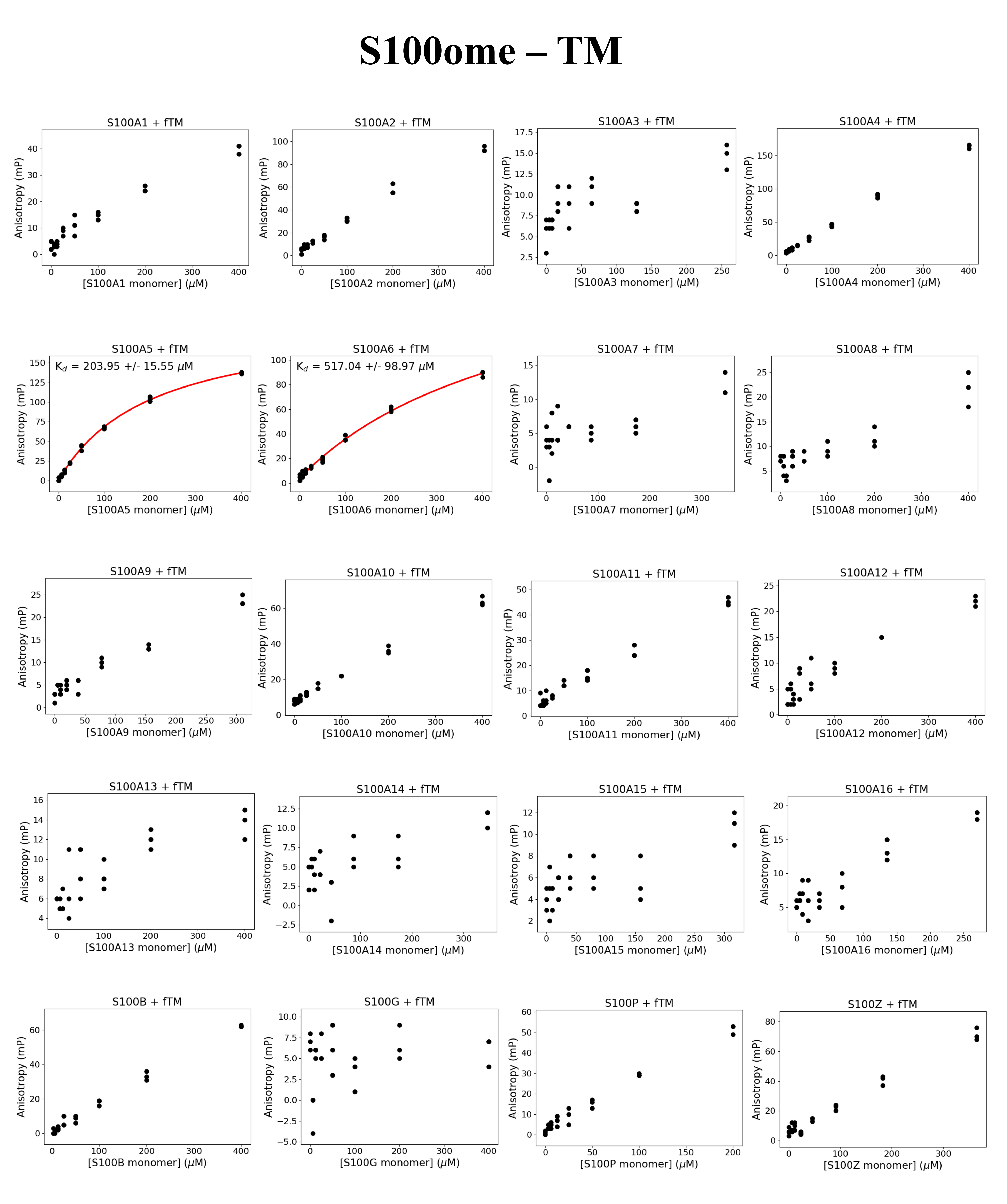

### Supplemental Fig 8

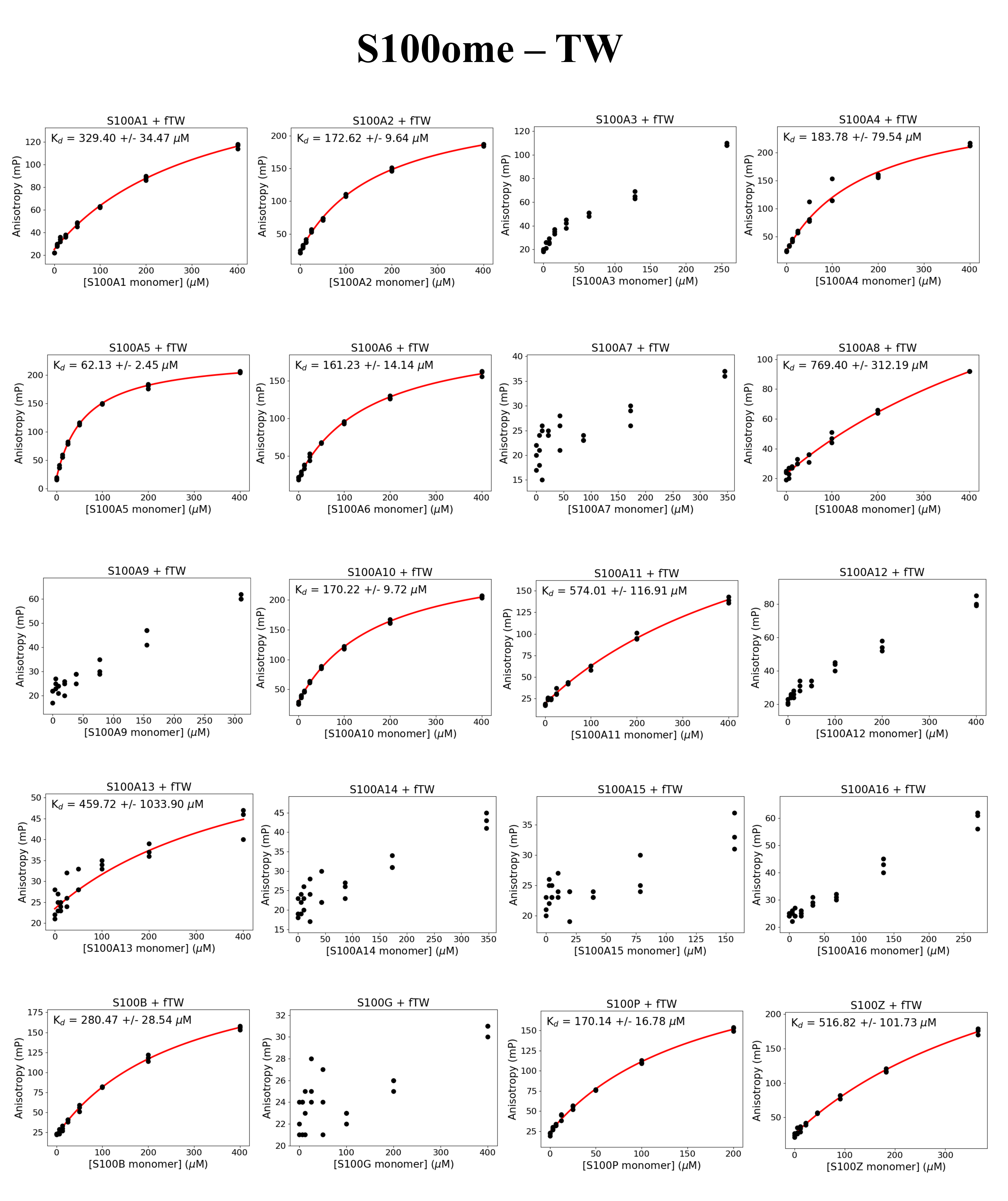

### Supplemental Fig 9

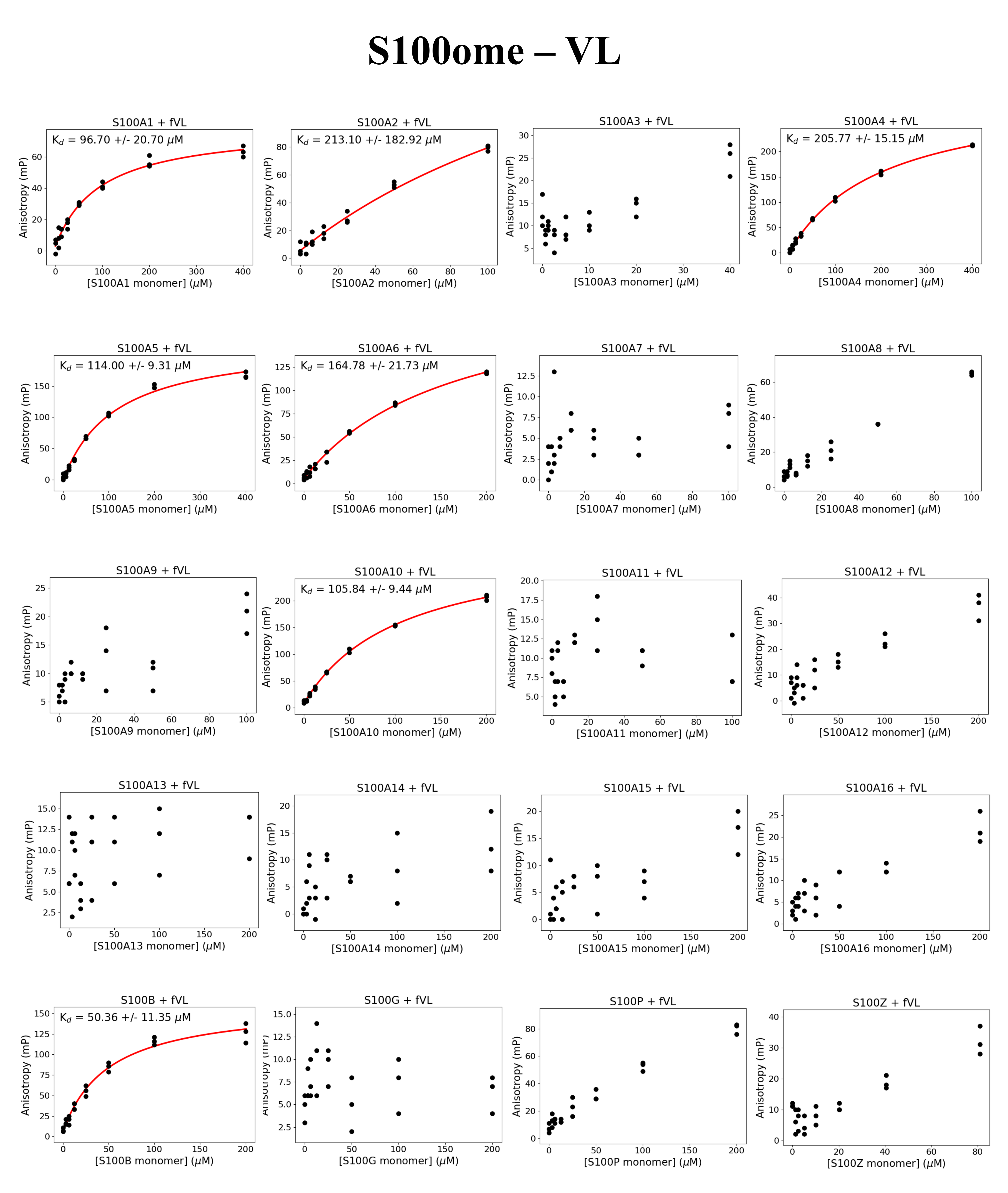

### Supplemental Fig 10

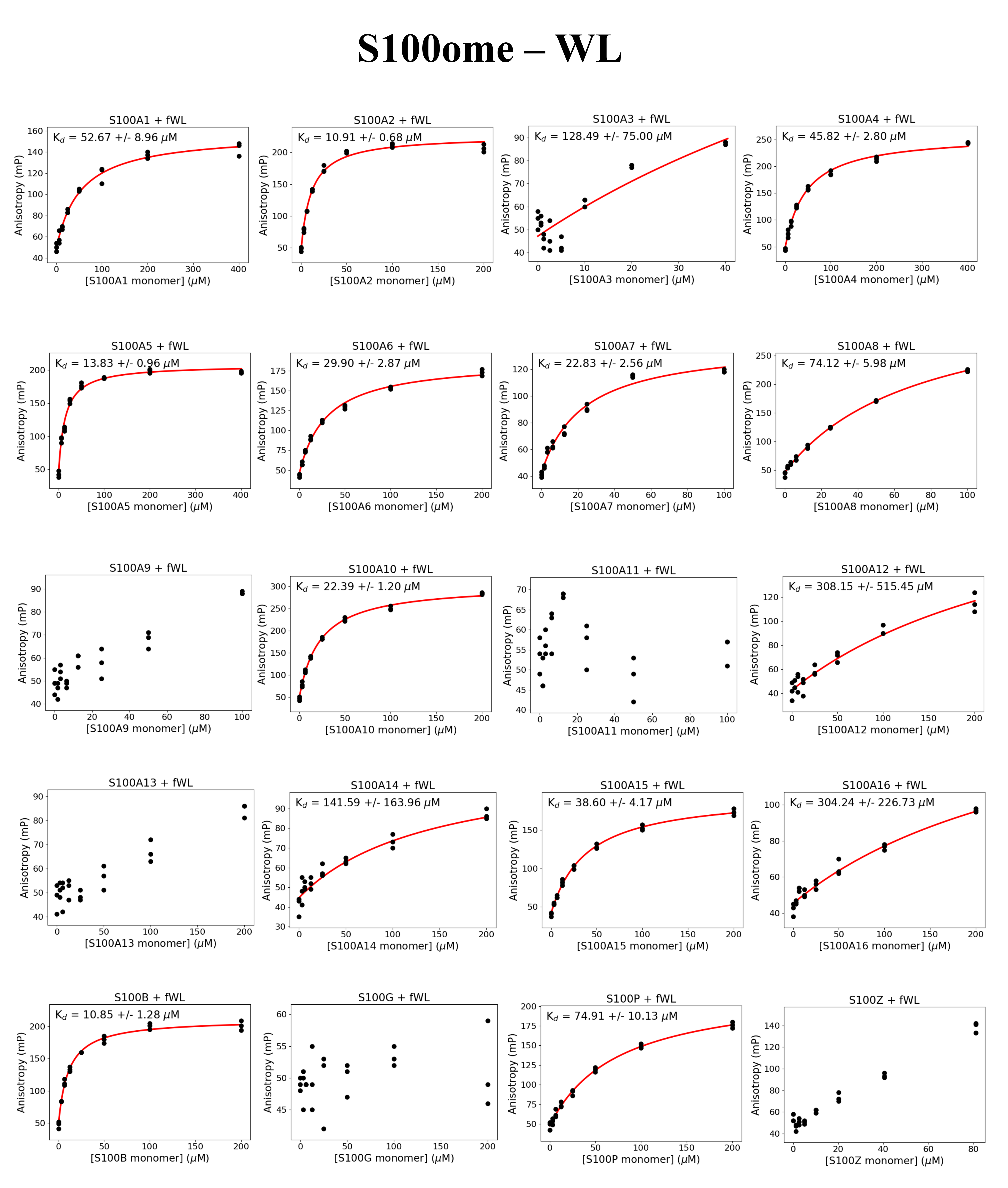

### Supplemental Fig 11

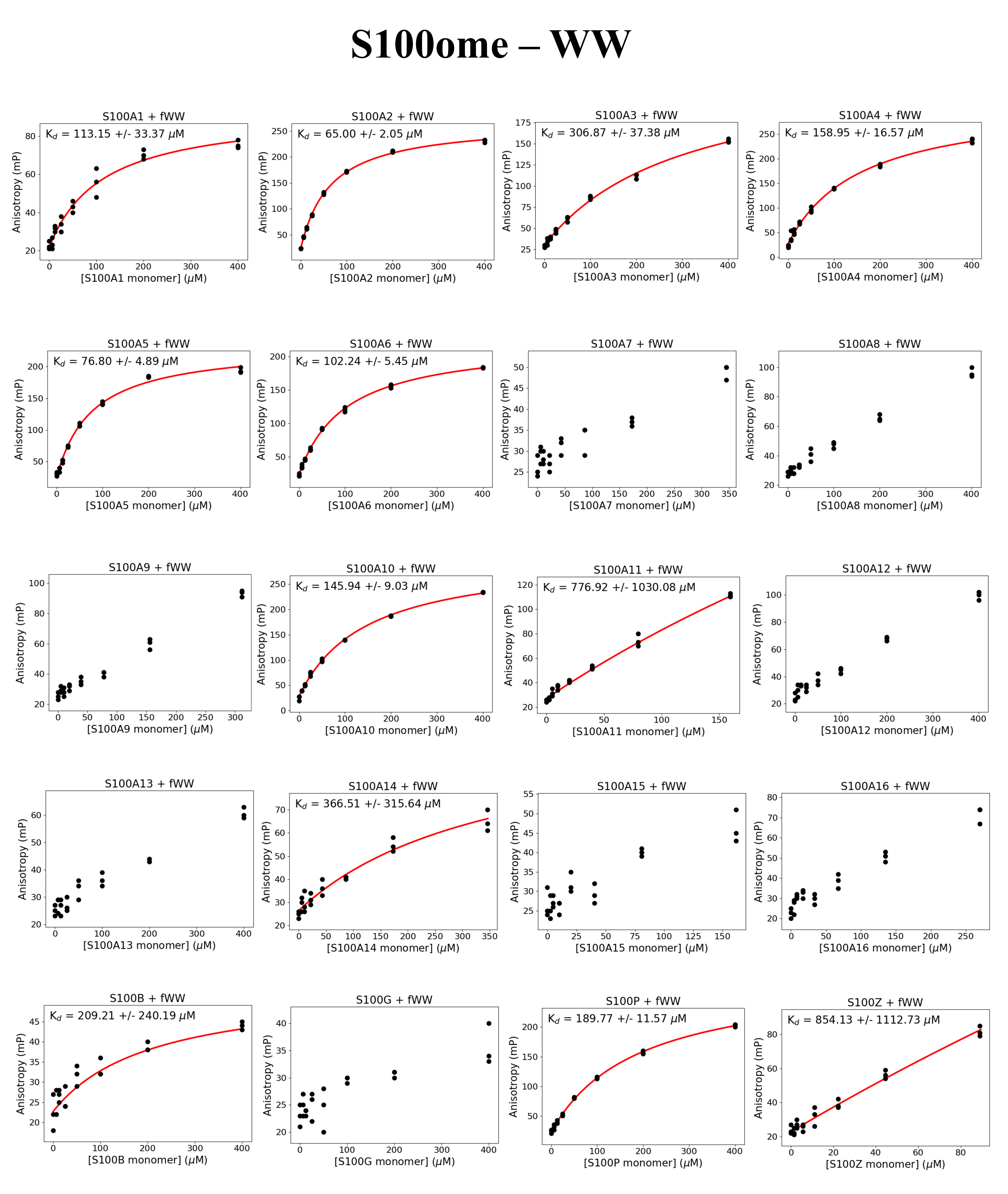

### Supplemental Fig 12

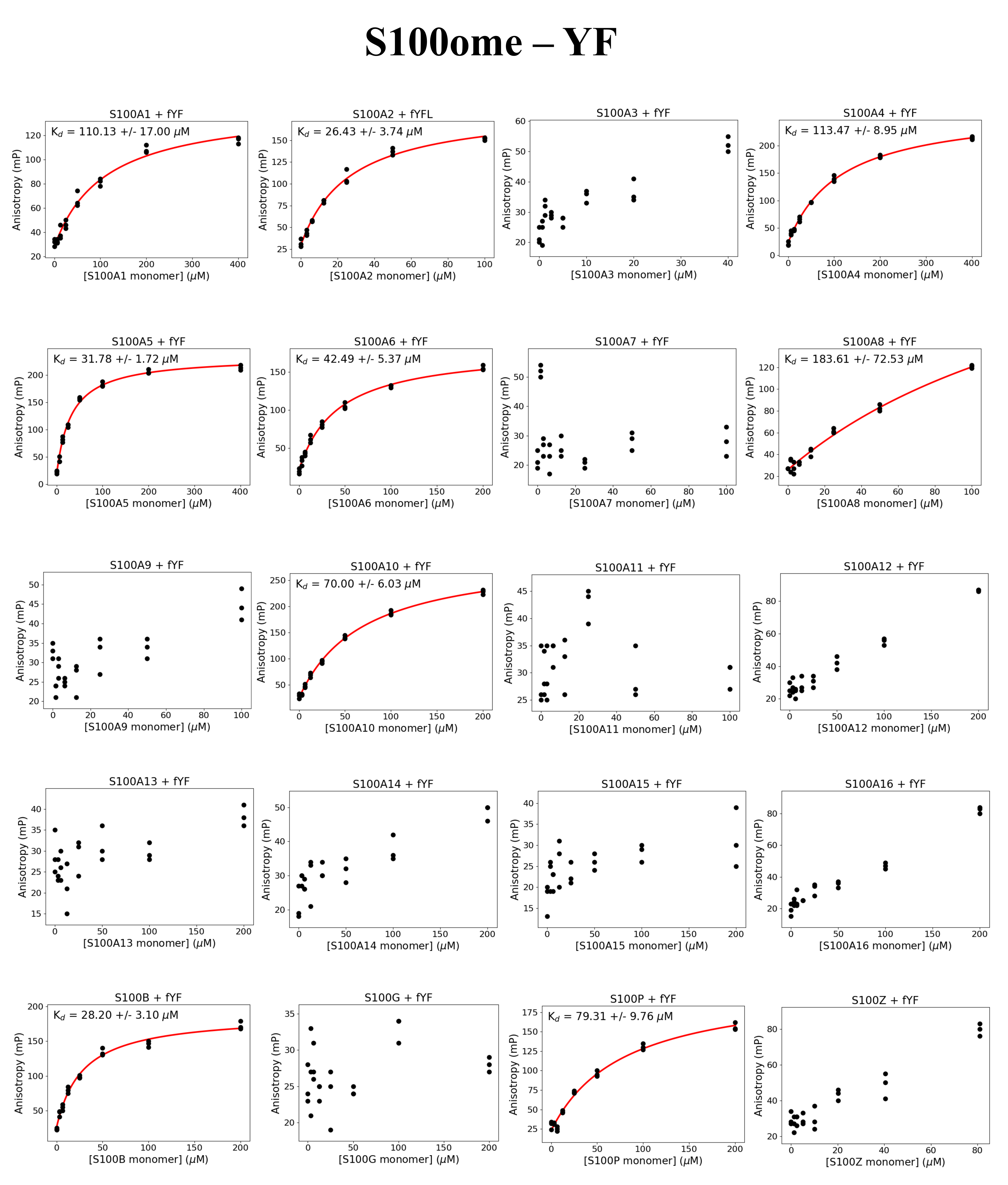

### Supplemental Fig 13

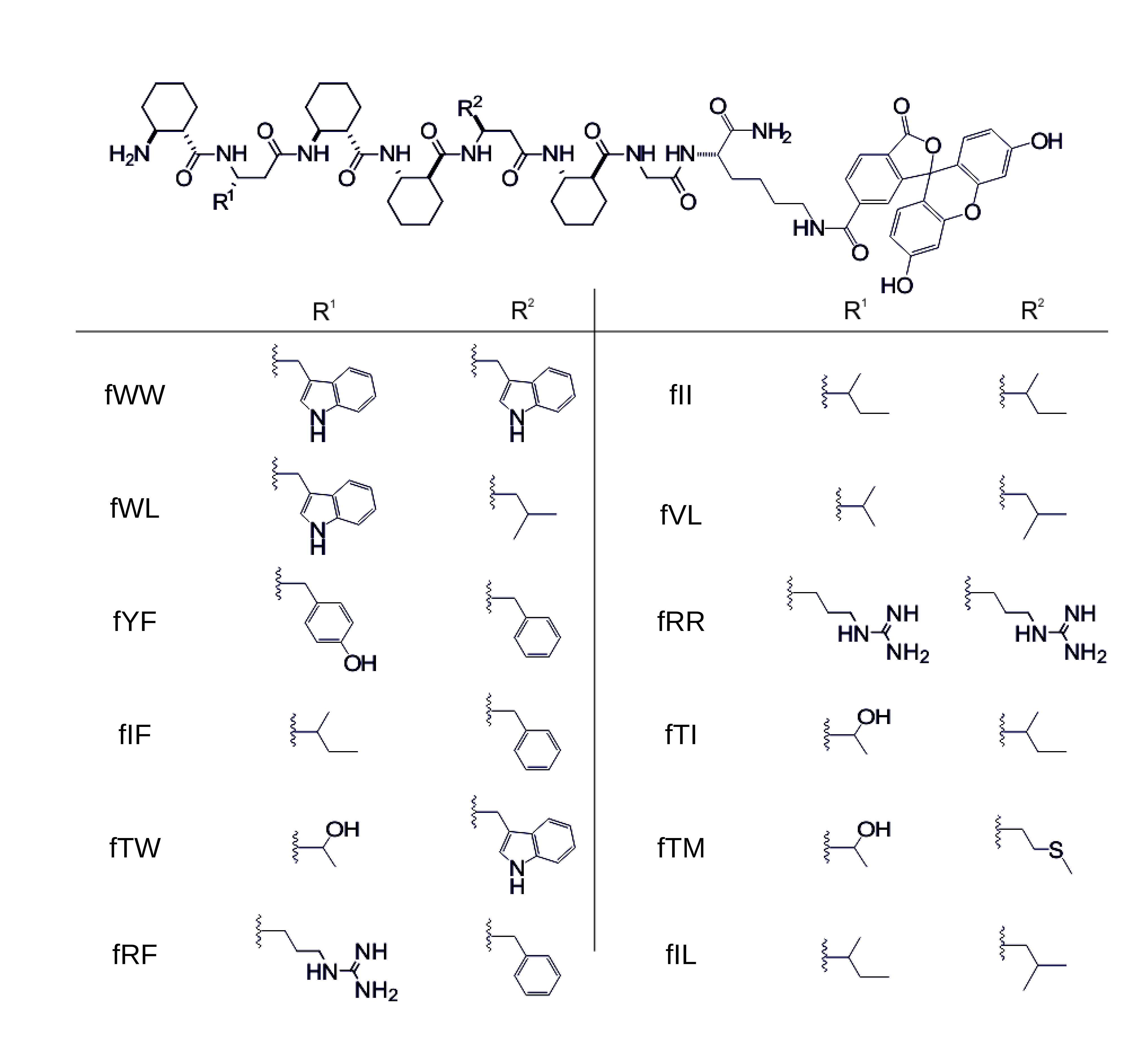

### Supplemental Fig 14

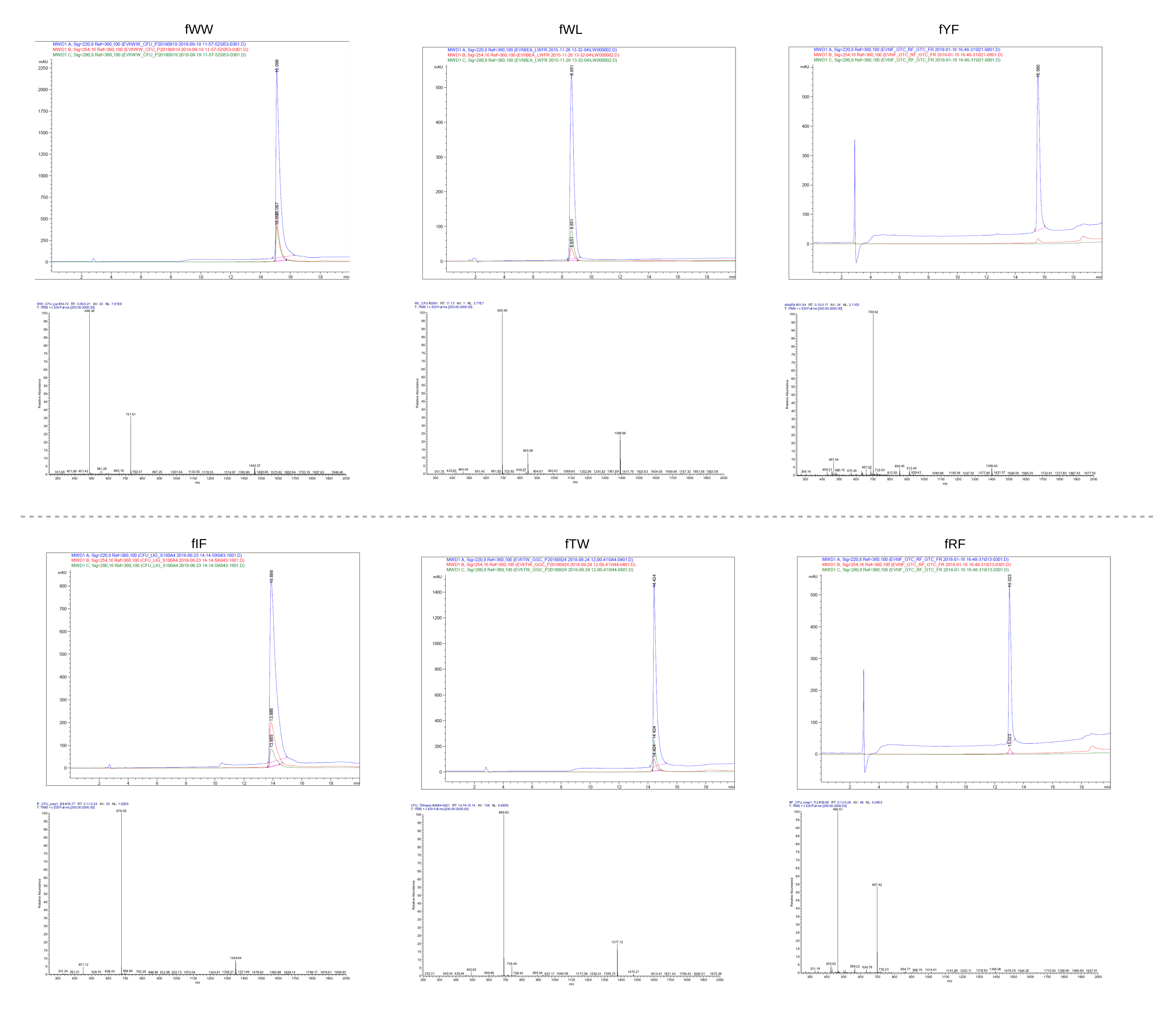

### Supplemental Fig 15

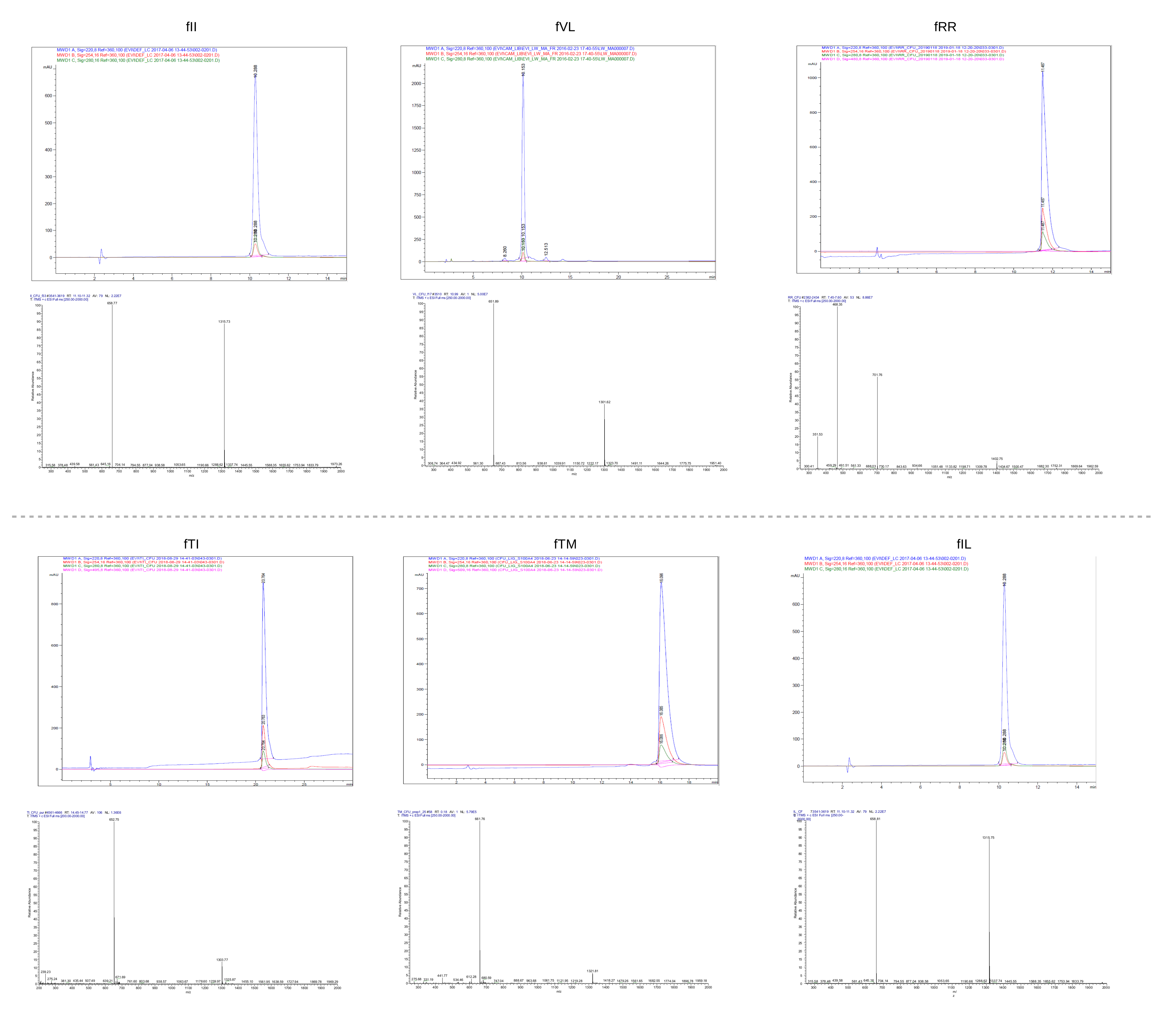

### Supplemental Table 1

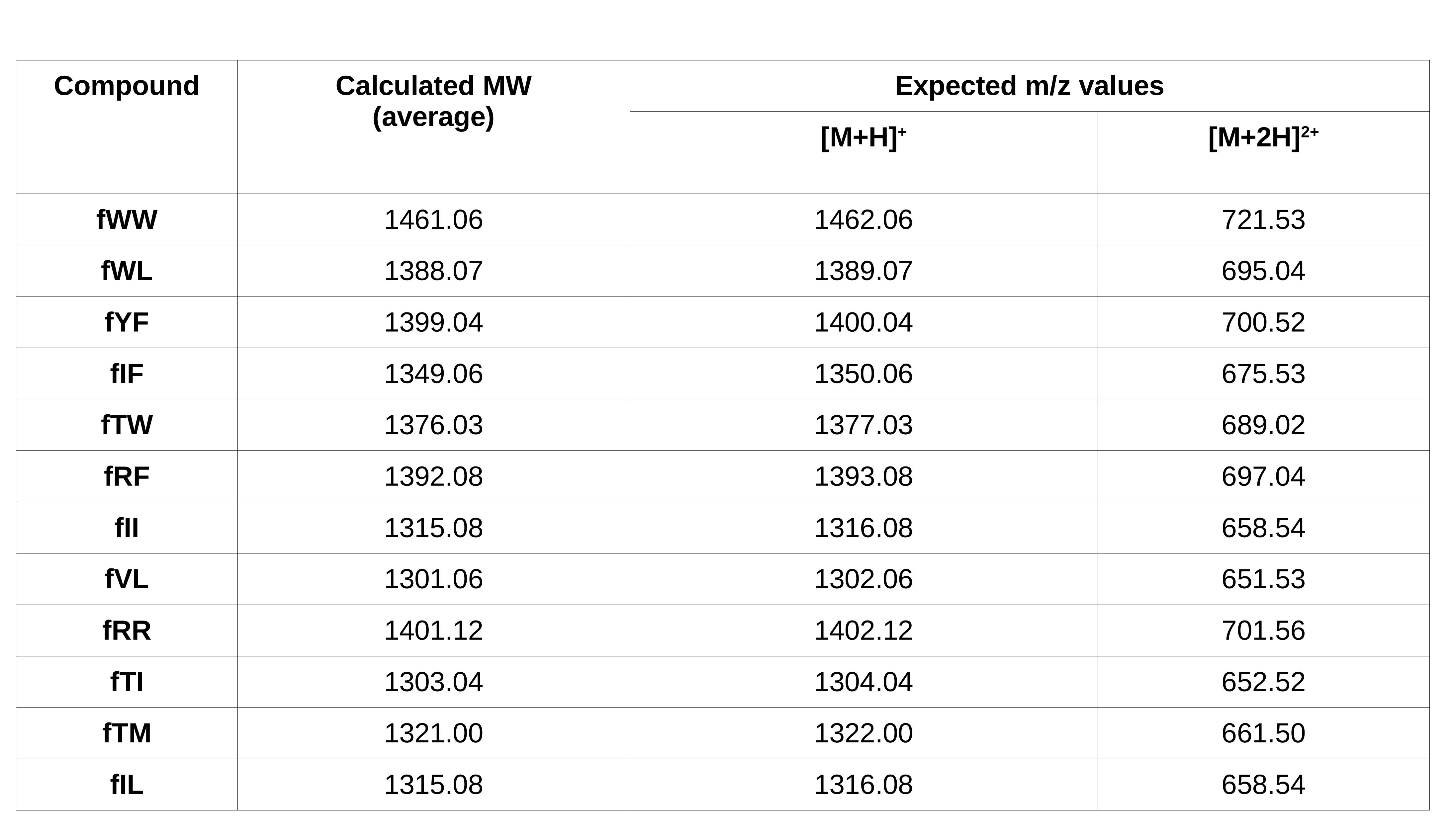
